# Cellular Stress Tolerance Governs Genetic Transformability in Recalcitrant *Candida* Species

**DOI:** 10.64898/2026.06.25.734625

**Authors:** Christopher J. Cotter, Dana L. Carper, Richard J. Giannone, Cong T. Trinh

## Abstract

*Candida* species are fungal pathogens whose rapidly increasing antifungal resistance poses a substantial public health challenge. High-throughput CRISPR-based screening could accelerate antifungal target discovery, yet its application in *Candida* has been limited by low DNA transformation efficiency. Chemical transformation exposes cells to environmental stresses to permit DNA uptake, but the physiological constraints on transformability remain poorly defined. Here, we show that genetic transformability in *C. albicans* is governed by the cellular capacity to withstand and recover from transformation-induced stress. Nutrient limitation markedly enhances transformation efficiency, while extracellular pH and lithium acetate chemistry strongly modulate this response. Systems-level proteomic analyses reveal that nutrient limitation and transformation chemistry prime oxidative stress tolerance, and transformation efficiency correlates with the expression of oxidative stress response proteins. Guided by these insights, we developed a generalizable fungal advanced chemical transformation (FACT) method that increases transformation efficiency across diverse *Candida* species and enables robust pooled CRISPR screening.

## INTRODUCTION

*Candida* species are opportunistic commensal yeasts that cause invasive fungal infections with elevated mortality rates ranging from 10-47%, depending on the infecting species, treatment, and patient-related factors such as age and comorbidities^1,2^. Current treatment options for invasive candidiasis remain limited to three major antifungal drug classes: azoles, echinocandins, and polyenes, and no new drug classes have been introduced in more than two decades^3^. While these drug classes have been historically effective, their widespread use in clinical and agricultural settings has accelerated the emergence of antifungal-resistant species, including the multidrug-resistant fungal pathogen, *C. auris*^4–7^. This combination of limited therapeutic options and rising resistance underscores the urgent need to identify novel drug targets and to improve our understanding of antifungal resistance mechanisms.

Among *Candida* species, *C. albicans* is both the most prevalent cause of invasive candidiasis and the most well-studied, making it a primary model for studying fungal pathogenesis and antifungal resistance^1,8,9^. However, despite decades of research, thousands of genes remain uncharacterized, largely due to technical challenges in genetic manipulation of *C. albicans,* including its diploid genome, lack of compatible plasmid systems, and reliance on integrative methods for genome modification^10–12^. These limitations have slowed the pace of functional genomics and hindered efforts to systematically discover new therapeutic targets at scale.

Pooled CRISPR screens provide a powerful framework for defining gene functions, identifying novel drug targets, and elucidating mechanisms underlying drug susceptibility in a high-throughput manner^13^. The development of CRISPR tools for some *Candida* species has expanded rapidly in recent years, enabling targeted gene disruptions and programmable transcriptional control^12,14–18^. However, the implementation of pooled CRISPR screens in genetically recalcitrant *Candida* species such as *C. albicans* is severely limited by low transformation efficiencies that restrict guide RNA (gRNA) library representation and overall screen robustness.

Chemical transformation is often the method of choice for transformation of *Candida* species due its simplicity and cost-effectiveness, yet even the most optimized protocols yield only 10^1^-10^2^ transformants per µg DNA in *C. albicans*^19^. Despite their practicality, these protocols rely on intense environmental stressors, including prolonged periods of starvation coupled with exposure to elevated temperatures and chemical treatments such as lithium acetate (LiOAc), ethylenediaminetetraacetic acid (EDTA), and polyethylene glycol (PEG), to transiently increase cell permeability and promote DNA uptake at the cost of substantial physiological stress that reduces viability^19–21^. As a result, chemical transformation represents a trade-off between DNA delivery and stress-induced cytotoxicity. Despite its central importance, how *Candida* cells tolerate, adapt to, or recover from transformation-associated stress remains poorly understood, constraining efforts to rationally improve transformation efficiency.

While distantly related, reports in *Schizosaccharomyces pombe* have demonstrated that preculturing cells in minimal media and adjusting the LiOAc pH can substantially improve chemical transformation efficiency^22^. Notably, nutrient limitation is well known to induce hormetic responses across diverse eukaryotes, shifting cells toward stress-adapted physiological states that favor survival and recovery^23,24^. These observations suggest that cellular physiological states with enhanced stress-tolerance capacity may be inherently more transformable. However, whether such stress-adaptive principles constrain chemical transformability in *Candida* species, and whether they can be systematically leveraged, has not been examined. We therefore hypothesized that modulating cellular stress physiology prior to and during transformation would enhance transformability in genetically recalcitrant *Candida* species.

Here, we demonstrate that chemical transformation efficiency is governed by cellular stress tolerance and recovery capacity rather than DNA uptake alone. By modulating culture and transformation chemistry, we develop a fungal advanced chemical transformation (FACT) method that significantly improves transformation efficiency across a diverse range of *Candida* species. Through physiological and proteomic analyses, we identify stress-associated metabolic burdens imposed by conventional transformation conditions and show that the FACT method promotes cellular states with enhanced stress tolerance, with oxidative stress resilience strongly correlating with transformation success. Finally, we demonstrate that the FACT method enables robust pooled CRISPR screening in *C. albicans*. These results establish a physiological framework for improving transformability and provide an enabling platform for large-scale functional genomics and drug target discovery in clinically important *Candida* species.

## RESULTS

### Nutrient Limitation Promotes Genetic Transformability in *C. albicans*

To determine whether physiological states influence transformability, we first examined how mild nutrient limitation during preculture affects transformation efficiency in *C. albicans*. Transformation efficiency was assessed using a Cas9-gRNA plasmid carrying our previously developed CRISPR-GRIT *ADE2* targeting cassette, which integrates at the *ENO1* locus^16^. Because CRISPR-induced DNA damage can reduce cell viability, the use of an active guide better reflects the requirements of pooled CRISPR screens. The CRISPR-GRIT design ensures coupled delivery of each gRNA with its corresponding repair template, while *ADE2* targeting provides a simple phenotypic readout of successful editing via red colony formation.

Using an established *C. albicans* chemical transformation protocol^19^ as the baseline (standard method), cells were cultured in either nutrient rich media (YPD) or minimal media (YNB) to prior to transformation. Preculturing cells in YNB rather than YPD resulted in a 10-fold increase in transformation efficiency (Figure 1A). Supplementing YNB with potassium acetate (KOAc), shown to improve transformation in *S. pombe*, further increased the transformation efficiency to 2.89 × 10^2^ per µg DNA (Figure 1A). Subsequent experiments were conducted using YNB with KOAc, hereafter referred to as YNB for simplicity.

**Figure 1.**
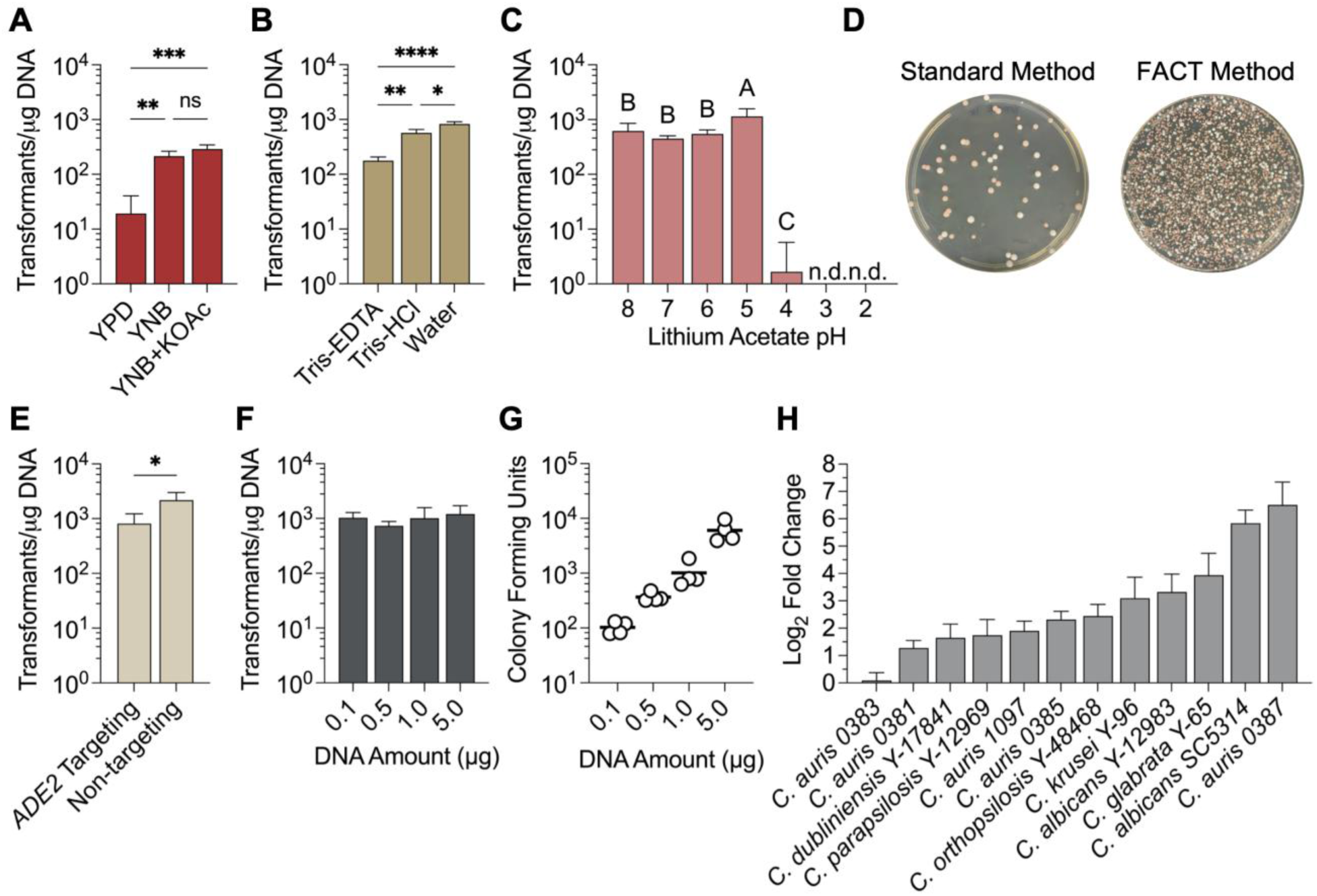
Preculture media and transformation chemistry influence transformation efficiency in *C. albicans.* **(A)** The effect of preculture media on *C. albicans* transformation efficiency using the linearized CRISPR-GRIT *ADE2* targeting DNA cassette. **(B)** The effect of the LiOAc solvent on *C. albicans* transformation efficiency after preculturing the cells in YNB+KOAc. **(C)** The effect of LiOAc pH on *C. albicans* transformation efficiency after preculturing the cells in YNB+KOAc. **(D)** Plate pictures of transformations of 5 µg of linearized CRISRP-GRIT *ADE2* using the standard method (left) and the FACT method (right). **(E)** Transformation efficiencies using different amounts of transforming DNA. **(F)** Colony counts (CFU) using different amounts of transforming DNA. **(G)** Log2 fold change improvement in transformation efficiency between the FACT and standard methods for different *Candida* species. Statistical significance was assessed using one-way ANOVA. For compact letter display, groups sharing a letter are not significantly different, whereas groups with different letters are. Data are the mean ± standard deviation of at least three biological replicates. *P < 0.05, **P < 0.01, ***P < 0.001, ****P < 0.0001.

### Transformation Chemistry Strongly Influences Genetic Transformability

To determine whether transformation chemistry further constrains transformability, we next examined how LiOAc formulation influences transformation efficiency in *C. albicans.* EDTA is commonly added to yeast transformation mixtures to destabilize the cell wall and cell membrane and protect DNA from nuclease degradation by chelating divalent cations; however, it is fungistatic in *C. albicans*^25–28^. Holding the pH constant at 8, we observed a 2.2-fold increase in transformation efficiency by removing the EDTA from the Tris-HCl buffer and a 3.7-fold increase by changing the LiOAc solvent to water (Figure 1B).

Using water as the LiOAc solvent, we examined the effect of the LiOAc pH on transformation efficiency, as pH strongly influences cellular physiology. In cells precultured in YNB, transformation efficiencies increased with decreasing LiOAc pH to an optimum at pH 5. Under these optimized conditions, which we refer to as the FACT method, transformations with the CRISPR-GRIT *ADE2* cassette routinely resulted in ∼10^3^ transformants per µg DNA (Figure 1C). The distinct difference between the standard and FACT methods can be visualized on solid plates (Figure 1D). Transformations using LiOAc with a pH below 5 resulted in little to no colonies due to a pronounced loss of cell viability (Figure S1A-B). Lowering the pH to 5 also increased the transformation efficiency of cells precultured in YPD (Figure S1C).

Using constructs lacking an active gRNA, transformation efficiencies were approximately 2-fold higher, due to the absence of Cas9-induced DNA damage (Figure 1E). Typical *C. albicans* transformation protocols require 1.5-10 µg of DNA for successful integration, a resource-intensive demand that can be impractical for large or low-copy plasmids, such as those containing Cas9^14,19,20^. Using the FACT protocol, transformants could be obtained with as little as 0.1 µg of DNA, with the transformation efficiency remaining constant across a range of 0.1-5 µg DNA (Figure 1F). Moreover, the number of colonies per transformation scaled linearly with the DNA amount (Figure 1G).

### Stress-Tolerant Transformation Is Conserved Across Diverse *Candida* Species

Non-*albicans Candida* species represent a diverse group of emerging fungal pathogens that are poorly characterized and genetically recalcitrant. Applying the FACT method to 10 non-*albicans* strains and one *C. albicans* clinicals increased transformation efficiency for all species tested relative to the standard method (Figures 1H, S2), with *C. krusei* and *C. auris* AR-0387 exhibiting the largest gains. As stress tolerance varies significantly among strains and species, strain specific tuning, including LiOAc pH, heat shock parameters, and recovery times, may further improve transformation efficiencies.

### Nutrient Limitation and Transformation Chemistry Enhance Cellular Stress Tolerance

#### FACT Conditions Accelerate Post-Transformation Recovery

Throughout the experiments, we noticed larger, more compact cell pellets following the recovery step for cells transformed using the FACT method compared to the standard method (Figure 2A), suggesting differences in post-transformation growth dynamics. We therefore hypothesized that the increased transformation efficiency under FACT conditions reflects improved tolerance to transformation-associated stress, leading to faster recovery, increased viability, or both.

**Figure 2.**
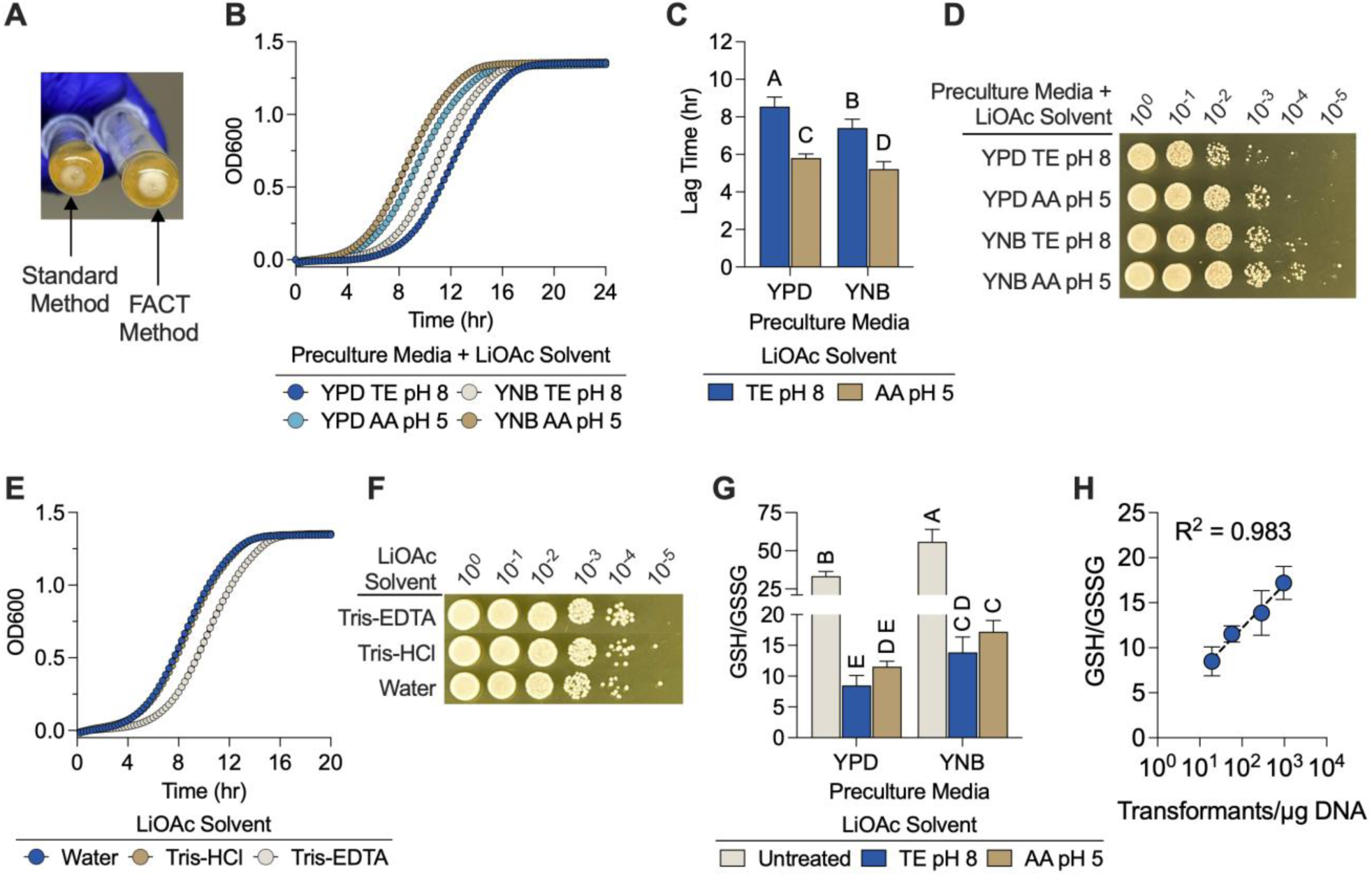
FACT conditions accelerate recovery, improve viability, and reduce oxidative stress, promoting transformation. **(A)** Cell pellets following recovery using the standard and FACT methods. **(B)** Post-heat shock recovery growth curves of *C. albicans* cells precultured in YPD or YNB and treated with 100 mM LiOAc and 50% w/v PEG 3350 dissolved in water titrated to pH 5 (AA pH 5) or in TE buffer at pH 8 (TE pH 8). **(C)** Lag time during the recovery growth for each treated sample. **(D)** Spotting viability assay following LiOAc/PEG treatment and heat shock. **(E)** The effect of LiOAc solvents on recovery growth kinetics for cells precultured in YNB. **(F)** Spotting viability assay for cells precultured in YNB and transformed using different LiOAc solvents. **(G)** Reduced to oxidized glutathione ratios (GSH/GSSG) for untreated and treated samples following heat shock. **(H)** Linear correlation between transformation efficiency and GSH/GSSG ratios. Statistical significance was assessed using two-way ANOVA followed by Tukey’s multiple comparisons test and represented with compact letter display: groups sharing a letter are not significantly different, whereas groups with different letters are. Data are the mean ± standard deviation of at least three biological replicates.

To test this, we quantified recovery kinetics following transformation. Independent of preculture media, cells treated with LiOAc and PEG at pH 5 (LiOAc pH 5), as implemented in the FACT method, exhibited a reduced lag phase, relative to cells treated with LiOAc and PEG in TE buffer (LiOAc TE) (Figure 2B-C). Holding the pH constant at 8 while varying the LiOAc solvent, cells treated with LiOAc TE displayed a prolonged recovery lag compared to those treated with LiOAc dissolved in either Tris-HCl or water (Figure 2E), suggesting that EDTA exacerbates physiological burden during recovery. Among cells treated with LiOAc TE, those precultured in YNB recovered more rapidly than cells precultured in YPD (Figure 2B-C), indicating that both preculture state and transformation chemistry contribute to recovery capacity.

#### FACT Method Refinements Increase Cell Viability

In parallel, we assessed post-transformation viability using spotting assays. Preculturing cells in YNB resulted in approximately an order of magnitude increase in viable cell counts relative to YPD-grown cells (Figure 2D). For YPD precultured cells, treatment with LiOAc pH 5 increased viability compared to LiOAc TE; however, this effect was less pronounced in YNB-precultured cells (Figure 2D), consistent with nutrient limitation establishing a more stress-tolerant state prior to transformation. Notably, we observed little difference in viability between LiOAc solvents (Figure 2F), suggesting that the EDTA-associated recovery lag is not due to overt cytotoxicity.

#### Nutrient Limitation and LiOAc Improvements in FACT Reduce Oxidative Stress

To further probe the physiological basis of improved recovery and viability, we measured the reduced to oxidized glutathione ratios (GSH/GSSG) before and after transformation as an indicator of cellular redox state, with higher ratios indicating less oxidative stress^29,30^.

Before transformation, cells precultured in YNB had significantly higher GSH/GSSG ratios than YPD-grown cells (Figure 2G). Following transformation, GSH/GSSG ratios decreased across all conditions, confirming that chemical transformation induces substantial oxidative stress. However, YNB-precultured cells retained higher GSH/GSSG ratios than YPD-grown cells after treatment. In addition, cells transformed using LiOAc pH 5 had higher GSH/GSSG ratios than those transformed with LiOAc TE. Collectively, these results indicate that both the preculture media and transformation chemistry modulate oxidative stress tolerance, and that lower oxidative burdens are strongly associated with higher transformation efficiency (Figure 2H).

The reduced GSH/GSSG ratios observed under TE buffer conditions suggest a potential inhibitory effect of EDTA on antioxidant enzyme function, possibly through chelation of divalent metal cofactors required by enzymes such as superoxide dismutase and catalase^31,32^. Consistent with this interpretation, measurements of relative hydrogen peroxide (H_2_O_2_) levels in the LiOAc/PEG transformation mixtures revealed elevated H_2_O_2_ levels in samples treated with LiOAc TE compared to LiOAc pH 5, irrespective of preculture conditions (Figure S3). In contrast, H_2_O_2_ was undetectable in samples treated with LiOAc pH 5, supporting the hypothesis that EDTA containing buffers may exacerbate transformation-induced oxidative stress.

### Proteome Remodeling Under Nutrient Limitation Primes Cells for Transformation Stress Tolerance

#### Minimal Media Shifts Proteome Resources from Translation to Energy Production, Amino Acid Biosynthesis, and Protein Turnover

To define the cellular basis for the transformation improvements using the FACT method, we examined the proteomes of *C. albicans* precultured in both YPD and YNB before treatment and after exposure to LiOAc/PEG formulated either in TE buffer or at pH 5 in water, immediately prior to heat shock (Figure 3A). This design isolates the proteomic response to chemical exposure from the confounding effects of thermal stress while preserving comparability across conditions. Principal component analysis (PCA) showed that preculture media was the primary source of variance (PC1, 19.6%), with transformation treatment contributing a secondary separation largely independent of LiOAc solvent (PC2, 15.1%; Figure 3B). Within each preculture medium, treatment conditions produced distinct proteomic profiles, a pattern corroborated by pairwise Pearson correlation (Figure S4A-C).

**Figure 3:**
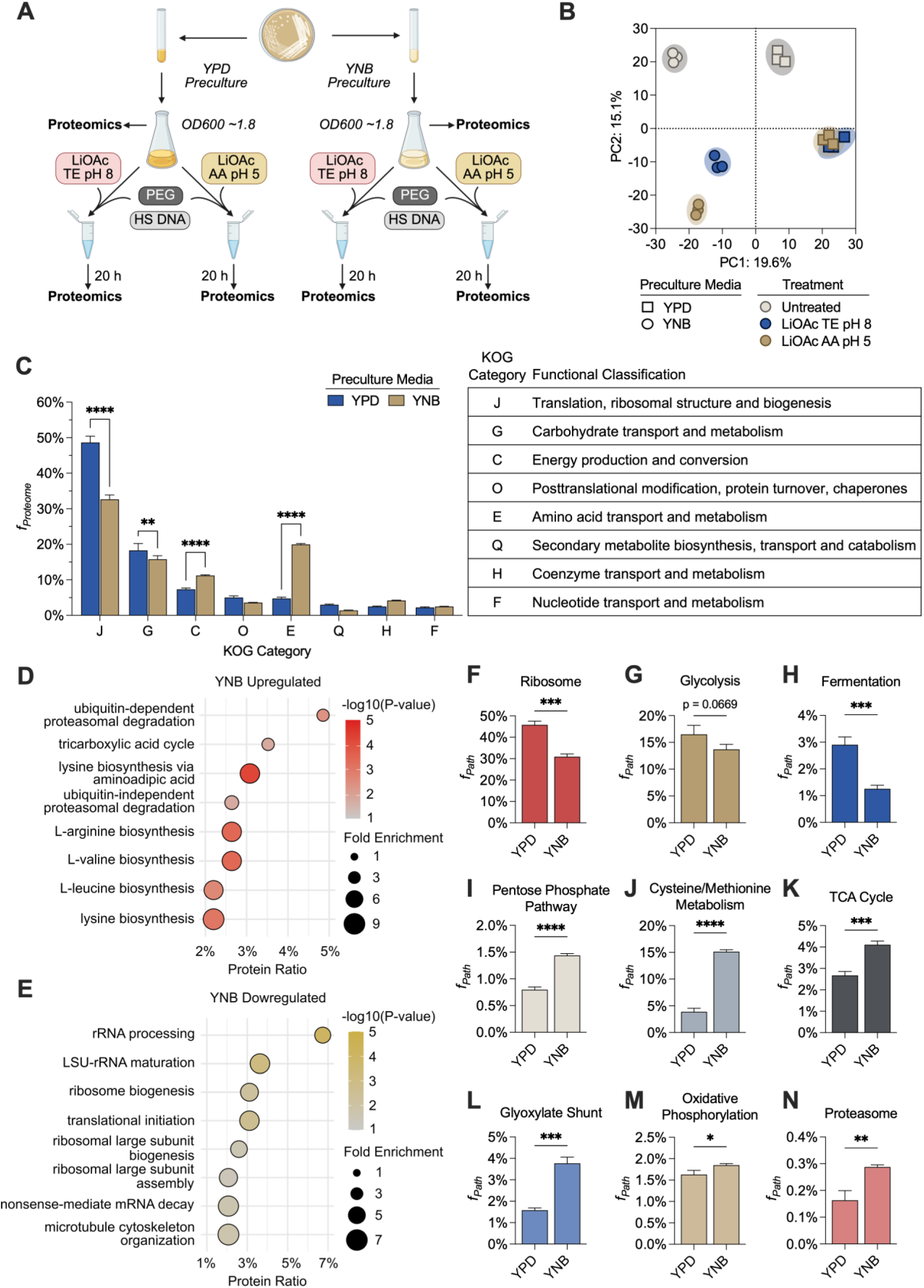
Minimal media primes cells for transformation stress tolerance by reallocating resources towards energy efficiency and redox homeostasis. **(A)** Proteomics sampling workflow. **(B)** PCA across all samples. **(C)** Allocation of proteome resources (*f_Proteome_*) devoted to top KOG categories between cells grown in YPD or YNB. Statistical significance was evaluated using two-way ANOVA. Multiple comparisons were corrected for using the Šídák correction. **(D-E)** Gene ontology biological process (GO BP) enrichment for proteins **(D)** upregulated or **(E)** downregulated in YNB relative to YPD. **(F-N)** Pathway proteome allocation for untreated cells grown in YPD and YNB. Statistical significance was evaluated using two-tailed unpaired t-tests. Data are the mean ± standard deviation of three biological replicates. *P < 0.05, **P < 0.01, ***P < 0.001, ****P < 0.0001.

Eukaryotic Orthologous Group (KOG) analysis revealed that, among untreated samples, YNB-grown cells redistributed proteome resources away from translation (KOG J) and carbohydrate metabolism (KOG G) and toward energy production (KOG C) and amino acid biosynthesis (KOG E) relative to YPD-cultured cells (Figure 3C). Functional enrichment of differentially expressed proteins corroborated this reallocation, identifying amino acid biosynthesis and tricarboxylic acid cycle (TCA cycle) proteins in the upregulated set (Figure 3D) and translation-associated proteins in the downregulated set (Figure 3E). Gene Ontology terms related to proteasome-mediated protein catabolism were also enriched among upregulated proteins (Figure 3D), a trend supported by increased proteome allocation to proteasome subunits in YNB-grown cells (Figure 3N). Priming proteasome activity by growing cells in nutrient-deplete media may enhance their ability to recycle resources, remove damaged components, and tolerate the stresses imposed during transformation^33^.

Quantitative pathway-level analysis further indicated extensive proteome rewiring of YNB-grown cells. Ribosomal investment was markedly reduced, while central carbon metabolism was redirected away from glycolysis toward the pentose phosphate pathway and respiratory ATP production via the TCA cycle and oxidative phosphorylation (Figure 3F-M). Among anabolic pathways, proteome investment in cysteine and methionine metabolism increased substantially, accounting for ∼15% of the total proteome in minimal media (Figure 3J). Because cysteine and methionine biosynthesis is energetically demanding with high protein, ATP, and NADPH costs^34,35^, this increase likely drives the global resource reallocation toward pathways supporting redox homeostasis and efficient energy generation, including the pentose phosphate pathway and respiration. Increased cysteine/methionine metabolism also supplies precursors for glutathione production which together with enhanced NADPH generation may elevate antioxidant capacity prior to transformation, consistent with the higher GSH/GSSG ratios in YNB (Figure 2D).

### Transformation Under Optimized Conditions Drives Stress-Adaptive Proteome Remodeling

We next examined how proteomes were remodeled in response to transformation stress. Differential expression analysis revealed that YNB-precultured cells mounted broader proteomic responses to transformation than YPD-grown cells across both LiOAc conditions (Figure 4A), with limited overlap in differentially expressed proteins between treatments (Figure S4D, F). Proteins shared between treatments were largely associated with the same preculture media, reinforcing that the preculture media strongly influences the proteome response.

**Figure 4:**
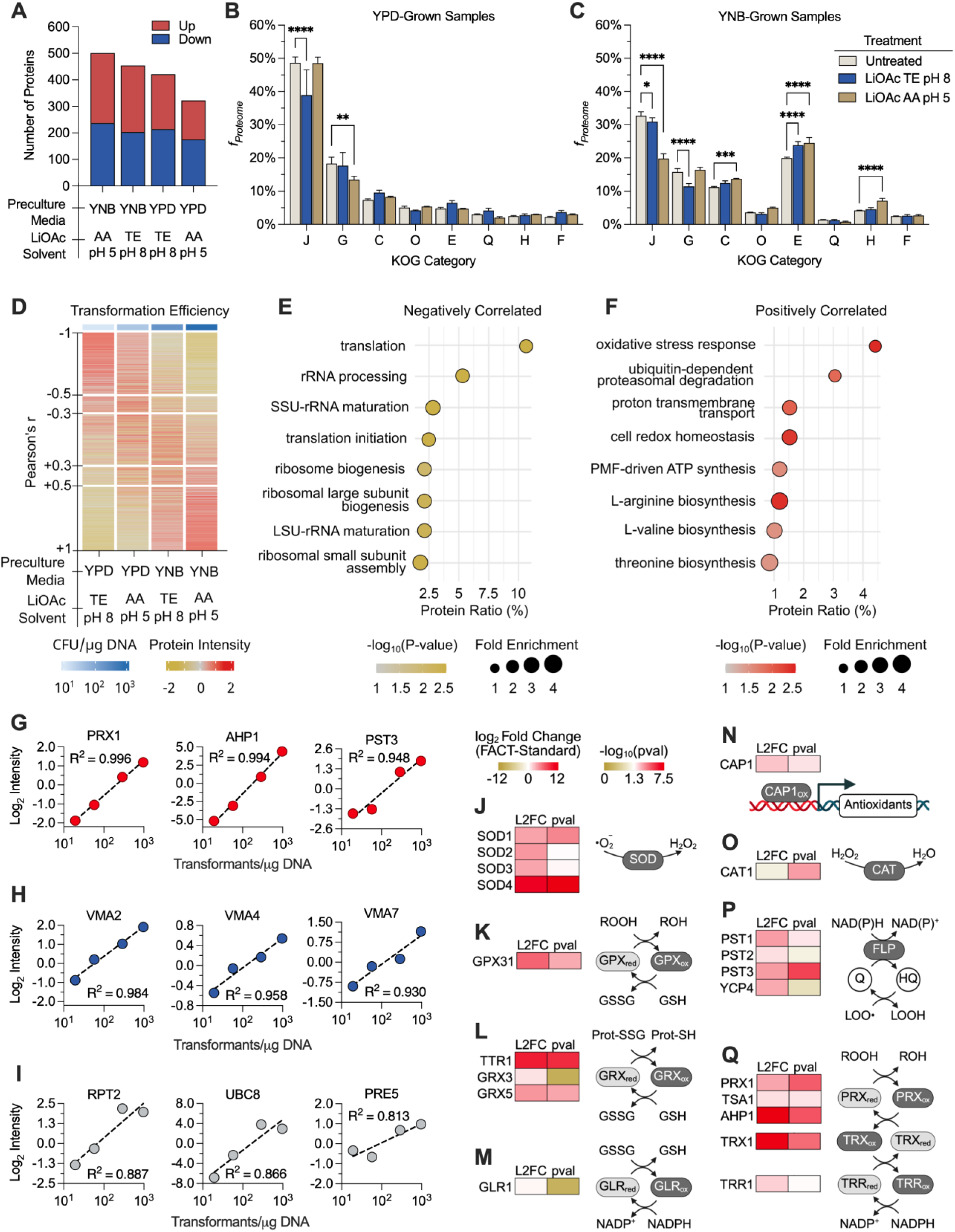
Proteome remodeling toward stress tolerance correlates with transformation efficiency. **(A)** Number of differentially expressed proteins identified for each treated sample relative the respective untreated sample (|log_2_ fold change| ≥ 1, adjusted p-value < 0.05). **(B-C)** Proteome allocation (*f_Proteome_*) devoted to top KOG categories between treated and untreated samples grown in either YPD **(B)** or YNB **(C)**. Statistical significance was evaluated using two-way ANOVA, followed by Tukey’s multiple comparisons test. **(D)** Pearson correlation between transformation efficiency (CFU/µg) and VSN protein intensity across all tested preculture and LiOAc conditions. **(E-F)** Gene ontology biological process (GO BP) enrichment for proteins that were **(E)** negatively correlated (r > 0.5) or **(F)** positively correlated (r < -0.5) with transformation efficiency. **(G-I)** Linear relationships for top positively correlated proteins involved in oxidative stress response **(G)**, proton transmembrane transport **(H)**, and proteasome-mediated ubiquitin-dependent catabolism **(I)**. **(J-Q)** Differential expression of oxidative stress response proteins between the FACT and standard methods. Categories include superoxide dismutases (SOD; **J**); glutathione system proteins including glutathione peroxidase (GPX; **K**), glutaredoxin (GRX; **L**), and glutathione reductase (GLR; **M**); the AP-1 transcription factor *CAP1* (**N**); catalase (CAT; **O**); flavodoxin-like proteins (FLP; **P**); and thioredoxin system proteins (**Q**), comprising peroxiredoxins (PRX), thioredoxin (TRX), and thioredoxin reductase (TRR). Abbreviations: LOO•, lipid peroxyl radical; LOOH, lipid hydroperoxide; ROOH, general peroxide (R = H or organic group); ROH, reduced peroxide; Q, quinone; HQ, reduced quinone (hydroquinone). P-values (pval) were calculated from moderated t-statistics using empirical Bayes shrinkage and adjusted for multiple testing by the Benjamini-Hochberg false discovery rate (FDR) method. Data are the mean ± standard deviation of three biological replicates. *P < 0.05, **P < 0.01, ***P < 0.001, ****P < 0.0001.

At the functional category level, transformation treatments induced proteome remodeling with heterogeneous changes across KOG classes (Figure S4E, G). Under FACT method conditions, cells exhibited a characteristic pattern of pronounced downregulation in amino acid biosynthesis and translation, coupled with strong upregulation in protein quality control (KOG O), indicating a strategic shift away from energy-intensive biosynthesis and toward proteostasis, hallmarks of adaptive stress responses in yeast^36,37^.

Global proteome allocation analyses reinforced these findings. YPD-grown cells exhibited limited proteomic remodeling after treatment, with only modest decreases in translation and carbohydrate metabolism (Figure 4B). Conversely, those cultured in YNB were able to significantly redistribute resources across multiple functional categories (Figure 4C). Following LiOAc pH 5 treatment, YNB-grown cells were able to substantially decrease their translation investment while increasing allocation to energy production, amino acid biosynthesis, and coenzyme metabolism (KOG H). Although many amino acid biosynthesis proteins were downregulated in DEP counts, the increased proteome fraction devoted to KOG E was driven mainly by high-cost methionine biosynthesis (Figure S5L). Proteome allocations toward other amino acid biosynthetic pathways were largely decreased or unchanged following transformation treatment, consistent with the general downregulation in biosynthetic capacity to adapt to stress (Figure S5). Together, these allocation changes show that transformation triggers broad stress-adaptive proteome reorganization, specifically under FACT conditions.

### Oxidative Stress Response Capacity Strongly Correlates with Transformability

To identify features associated with increased transformation efficiency, we correlated normalized protein abundances with transformation efficiencies across conditions (Figure 4D). Negatively correlated proteins (r < -0.5) were enriched for translation-associated functions (Figure 4E), reinforcing that high investment in growth-associated machineries characteristic of nutrient-rich states is detrimental to transformation. Positively correlated proteins (r > 0.5) were enriched for processes related to oxidative stress tolerance, redox homeostasis, proton transmembrane transport, and proteasome-mediated protein degradation (Figure 4F), including antioxidant enzymes (Figure 4G), V-ATPase subunits (Figure 4H) and proteasome-associated proteins (Figure 4I). These associations suggest that enhanced ROS detoxification and protein quality control capacities are defining features of highly transformable states, consistent with elevated GSH/GSSG ratios observed under FACT conditions.

#### Antioxidant Enzyme Expression is Elevated by the FACT Method

Direct comparison of antioxidant proteins between the FACT and standard methods showed broad upregulation of critical antioxidant proteins including peroxiredoxins (*PRX1, TSA1, AHP1*), thioredoxin and thioredoxin reductase (*TRX1, TRR1*), superoxide dismutases (*SOD1-4*), glutaredoxins and glutathione peroxidases (*TTR1, GRX5, GPX31*) and flavodoxin-like proteins (*PST1-4, YCP4*; Figure 4J-Q). Notably, *AHP1* (log_2_FC = 10.72, p. adj = 3.88 x 10^-5^), *TRX1* (log_2_FC = 10.43, p. adj = 1.20 x 10^-4^), *SOD4* (log_2_FC = 9.51, p. adj = 2.05 x 10^-7^), and *TTR1* (log_2_FC = 8.47, p. adj = 5.26 x 10^-6^) exhibited substantial increases relative to the standard method, indicating an enhanced capacity to detoxify diverse reactive oxygen species. Additionally, the AP-1 like transcription factor *CAP1,* a major transcription factor involved in oxidative stress tolerance, was significantly upregulated (log_2_FC = 1.99, p. adj = 1.62 x 10^-2^) using the FACT method^38^. These findings indicate that FACT places cells in a physiological state that supports a more effective oxidative stress response, which in turn aligns with the increased viability, faster recovery, and higher transformation efficiency observed.

### Inducing Oxidative Stress-Adapted States Enhances Transformability

Sublethal H_2_O_2_ can transiently establish an oxidative stress-adapted physiological state in yeast^38–43^. Building on the reduced oxidative burden and elevated stress-response capacity observed under FACT, we tested whether brief H_2_O_2_ preconditioning prior to transformation could phenocopy key features of this state and enhance transformability. Because *C. albicans* lacks a canonical general stress response and displays limited cross-protection between distinct stressors, H_2_O_2_ preconditioning is highly oxidative-stress specific rather than broadly protective^41^.

Cultures grown in YPD or YNB were exposed to 0.4 mM H_2_O_2_ one hour prior to collection (Figure 5A). Pretreated cells displayed increased tolerance to subsequent high-dose H_2_O_2_ exposure (Figure 5B). When transformed with a non-targeting CRISPR-Cas construct, H_2_O_2_ preconditioning increased transformation efficiency ∼2-fold (Figure 5C) and shortened recovery lag times by ∼20% (Figure 5D) in both media backgrounds, independent of LiOAc solvent. Cell viability was largely unchanged (Figure S6). These results suggest that mild oxidative preconditioning establishes a stress-adapted state that primarily accelerates post-transformation recovery, thereby improving transformation efficiency even under conditions that already confer baseline stress tolerance. This supports a model in which activation of oxidative stress response pathways improves transformability and suggests stress preconditioning as a broadly useful strategy for enhancing transformation efficiency.

**Figure 5:**
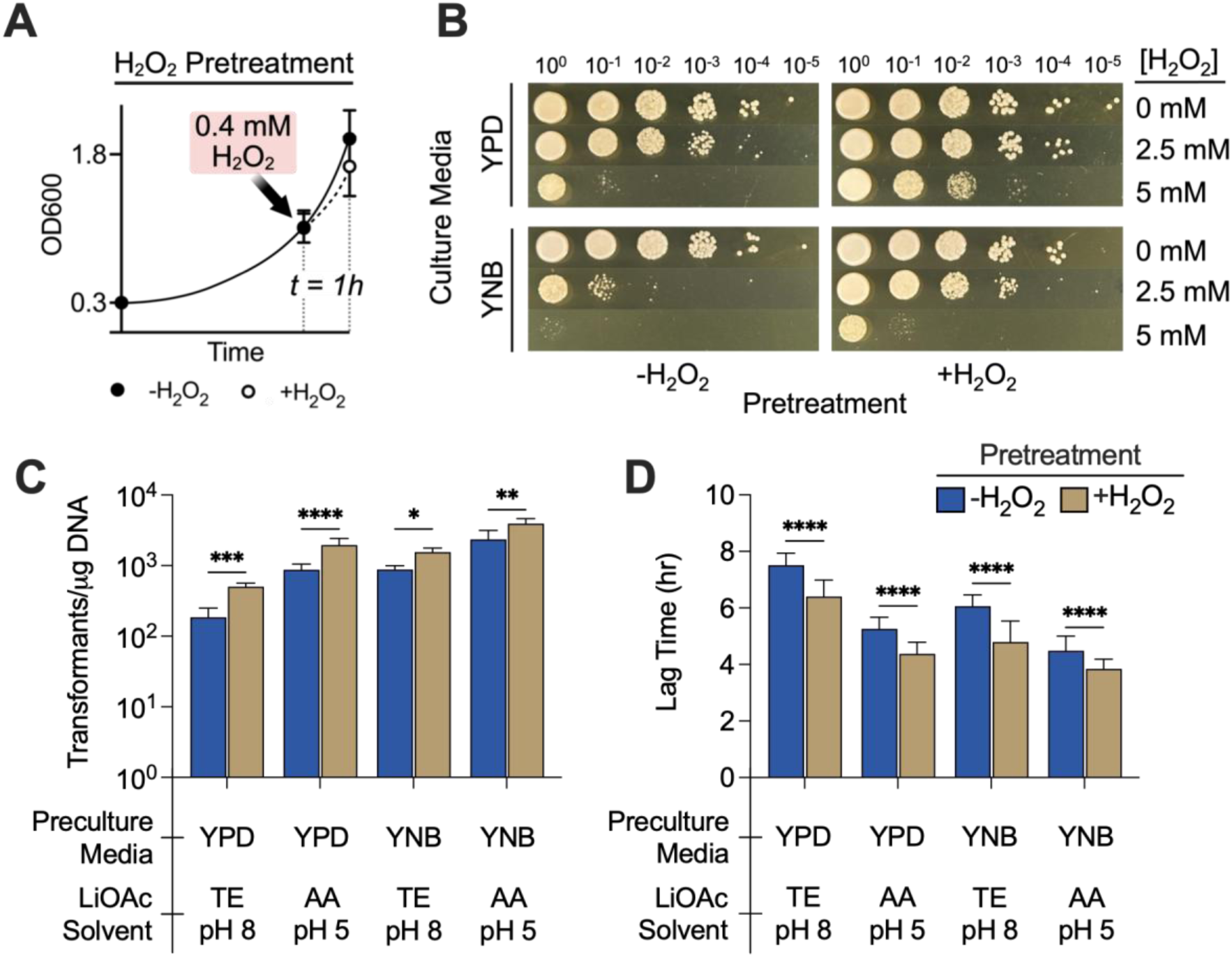
Oxidative stress priming enhances transformability. **(A)** Workflow of H_2_O_2_ treatment during preculture step. **(B)** Spotting assay assessing whether sublethal H_2_O_2_ preconditioning enhances H_2_O_2_ tolerance. YPD- or YNB-grown cells, with or without H_2_O_2_ pretreatment, were serially diluted and spotted onto YPD plates containing 0, 2.5, or 5 mM H_2_O_2._ Images were taken after 24 hours and are representative of three biological replicates. **(C-D)** Effect of H_2_O_2_ pretreatment on **(C)** transformation efficiency and **(D)** recovery lag time of cells grown in either YPD or YNB and transformed using LiOAc dissolved in either TE buffer at pH 8 (TE pH 8) or water titrated to pH 5 with acetic acid (AA pH 5). Statistical significance was evaluated using unpaired two-tailed t-tests. Data are the mean ± standard deviation of at least three biological replicates. *P < 0.05, **P < 0.01, ***P < 0.001, ****P < 0.0001.

### FACT Enables Robust Pooled CRISPR Screening in *C. albicans*

To assess the utility of the FACT method for high-throughput functional genomics, we performed a small-scale pooled gRNA library screen using 35 CRISPR-GRIT gRNAs targeting essential genes (*RPS15, TRL1, RPB2*), DNA repair genes (*RAD51, RAD52, LIG4*), biofilm-related genes (*ALS1, ACE2, FLO8*), three null guides, and previously utilized *ADE2* and *URA3* controls (Table S1). Initial samples (T0) were collected immediately following the 4-hour recovery period, and depleted samples were taken following a 40-hour selective outgrowth with nourseothricin (T40) (Figure 6A-B).

**Figure 6:**
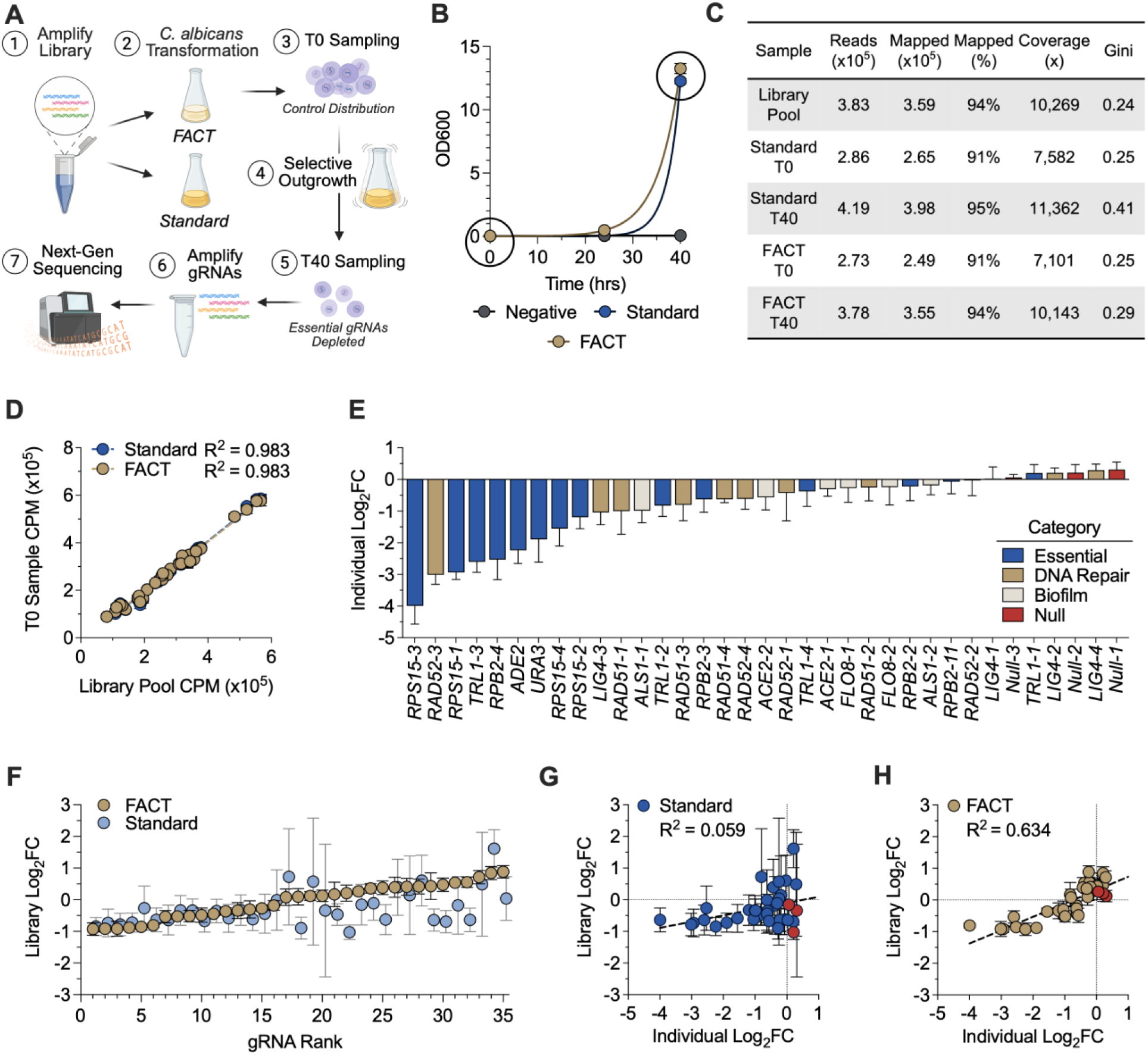
FACT method enables robust pooled CRISPR screens in *C. albicans*. **(A)** CRISPR screen workflow. **(B)** Growth curves during library depletion outgrowth for each condition. Samples were taken at t = 0 and t=40 as indicated by circled points. **(C)** Sequencing quality control metrics averaged for each sample. **(D)** Correlation between gRNA read counts (CPM, counts per million) detected in the pretransformed library pool and the initial (t=0) library samples. **(E)** Individual characterization of colony reduction for each CRISPR-GRIT gRNA in the pooled library relative to a non-targeting control **(F)** Library depletion values for each gRNA using the FACT and standard methods. **(G-H)** Correlations between individually characterized gRNA colony reductions and gRNA depletions in the library samples using the **(F)** standard transformation method and **(G)** FACT transformation method. Null gRNA data points are filled red. Data are the mean ± standard deviation of three biological replicates.

Sequenced gRNA loci from all samples exhibited good coverage, with >90% of reads mapped to gRNAs and ∼10^4^ reads counted per gRNA (Figure 6C). All gRNAs were detected in all samples, with no zero-count guides observed. Read counts from the T0 control samples for both transformation methods tightly correlated (R^2^=0.983) with those from the pretransformed library pool, suggesting little sequence loss during the transformation process (Figure 6D). Consistent with this, Gini index values were ∼0.25 for the pretransformed and T0 samples, with a modest increase in the depleted samples (Figure 6C). Given the small library size, this level of inequality is expected and indicative of acceptable gRNA representation rather than substantial bottlenecking or amplification bias.

To benchmark depletion accuracy, we individually characterized the fitness effects of each guide by quantifying colony reduction relative to a null control (Figure 6E). Essential gene gRNAs produced the strongest reduction in colony counts, whereas targeting DNA repair genes, biofilm genes, and null targets showed moderate to minimal effects (Figure 6E).

Each transformation method produced distinct gRNA depletion profiles (Figure 6F). When the pooled library depletion results were compared to the individually measured phenotypes, the standard transformation method performed poorly, yielding little correlation (R^2^=0.059) and high variability among replicates (Figure 6G). Additionally, the null gRNA controls were depleted. This likely reflects the low viability and stochastic, delayed recovery associated with the standard method, which can distort gRNA representation during selective outgrowth.

In contrast, the FACT method produced a substantially stronger correlation (R^2^=0.634) between the pooled depletion and the individual characterization, with markedly lower variance and expected control behavior (Figure 6H). These results demonstrate that the FACT method generates a more uniform and physiologically robust transformed population, enabling accurate gRNA representation through recovery and selective outgrowth. Together, these findings show that the FACT method overcomes a major barrier to high-throughput genetic screening in *C. albicans*, enabling reliable pooled CRISPR screens and expanding the tractability of this clinically important fungal pathogen for functional genomics.

## DISCUSSION

Efficient transformation is central to functional genomics, yet *Candida* species have remained among the most genetically recalcitrant fungal species for decades. In this study, we demonstrate that chemical transformation efficiency in *Candida* is governed not only by DNA uptake, but more fundamentally by the capacity of cells to tolerate and recover from transformation induced stress. By systematically modulating preculture conditions and transformation chemistry, we developed the FACT method, which improves transformation efficiency by more than two orders of magnitude across diverse *Candida* species and enables robust pooled CRISPR screens. Beyond a technical improvement, this work establishes physiological states associated with stress tolerance as a central determinant of transformability.

Chemical transformation exposes cells to multiple concurrent stressors, including nutrient deprivation, chemical insult, heat shock, and as demonstrated in this work, significant oxidative stress. Previous efforts to improve transformation efficiency have focused on intensifying select stressors to increase cell permeability for DNA uptake; however, these approaches yield only marginal gains while imposing severe burdens that reduce viability^19,21,44,45^. Our results instead demonstrate that transformability is an emergent property of the cellular state. Cells optimized for rapid growth are intrinsically poorly suited for chemical transformation, whereas cells preconditioned for stress tolerance, through nutrient limitation or oxidative priming, exhibit increased survival, faster recovery, and dramatically higher transformation efficiency.

Growth in rich medium commits cells to a physiological state optimized for rapid proliferation rather than stress tolerance. YPD-grown cells allocate a large fraction of their proteome to translation, ribosome biogenesis, and fermentative metabolism, maximizing growth rate at the expense of redox buffering, proteostasis capacity, and metabolic flexibility. While advantageous under permissive conditions, cells in this growth-optimized state exhibited a limited capacity to reprogram their proteomes to accommodate transformation stress, instead continuing to exhaust resources on energy intensive growth processes, leading to delayed recovery, reduced viability, and poor transformation efficiency.

In contrast, nutrient limitation primes *C. albicans* for subsequent transformation stress by globally reallocating proteome resources towards a physiological landscape more closely matched with the demands of chemical transformation. Reduced investment in translation lowers energetic burdens, while increased allocation toward respiration, the pentose phosphate pathway, cysteine and methionine metabolism, and proteasome function enhances energy efficiency, NADPH generation, redox buffering, and protein quality control, processes essential for surviving starvation, oxidative, and heat stress.

Consistent with this primed state, YNB-grown cells exhibited pronounced proteome flexibility following transformation stress, particularly under FACT conditions, further downregulating energy-intensive biosynthetic processes while upregulating protein turnover and oxidative stress response pathways. Notably, the abundance of oxidative stress response proteins showed a strong positive correlation with transformation efficiency and, together with our observed decrease in oxidative stress, supports enhanced redox tolerance as a defining feature of highly transformable states. This capacity for rapid proteome reprogramming toward oxidative stress tolerance enables YNB-grown cells to better withstand transformation-associated stress and efficiently resume growth following treatment, resulting in improved transformation outcomes.

Oxidative preconditioning provided an independent test of our stress-adapted physiological state model. Brief, sublethal H_2_O_2_ exposure, known to induce hormetic activation of the oxidative stress response^39–43^, prior to transformation increased transformation efficiency and reduced recovery lag. Importantly, because *C. albicans* lacks a canonical general stress response^41^, this improvement is unlikely to reflect broad stress tolerance or cross-protection; instead, it suggests that shifting cells into an oxidative stress-adapted program can increase transformation efficiency by enabling rapid recovery from transformation stress.

In addition to preculture state, the transformation chemistry itself influenced the overall stress burden. EDTA-containing LiOAc formulations increased oxidative burden and significantly prolonged recovery lag times. By removing EDTA and optimizing the LiOAc pH, the physiological burden imposed by transformation was reduced, leading to improved transformation efficiencies irrespective of initial culture conditions.

Notably, both recovery kinetics and viability emerged as a critical determinant of transformation success and are particularly important for pooled CRISPR screens, where stochastic losses during recovery can severely distort gRNA representation. Consistent with this, FACT enabled more reproducible CRISPR gRNA depletion profiles and yielded stronger correlations between pooled screen outcomes and individual gRNA phenotypes. These improvements highlight how physiological heterogeneity during transformation can propagate into large scale genetic screens, and how aligning transformation conditions with stress adapted cellular states directly improves screen accuracy.

Although developed for *Candida* species, the principles underlying FACT are likely generalizable. Stress preconditioning and hormetic responses are conserved strategies across microorganisms, and modulating cellular states to enhance transformation stress tolerance may increase viability and therefore transformation efficiency across kingdoms^23,46^. By establishing stress tolerance as a key determinant of transformation and developing a baseline method with improved tolerance, this work opens avenues to further enhance transformation efficiency through targeted manipulation of cellular physiology, including modulation of nutrient signaling, redox homeostasis, and stress-tolerance pathways.

Despite the substantial gains achieved here, transformation efficiencies remain lower than those observed in model yeasts, indicating that additional constraints persist. Cell wall architecture, endocytosis of DNA, and DNA integration mechanisms may further limit transformability and warrant future investigation. Additionally, development of stable plasmid systems that bypass the need for integration will greatly improve transformation efficiencies in *C. albicans*.

In summary, this study identifies oxidative stress tolerance and proteome flexibility as key drivers of chemical transformation efficiency in *Candida* species. By enhancing cellular stress tolerance, the FACT method provides both a practical tool and a conceptual framework for enabling high-throughput functional genomics in genetically recalcitrant fungal pathogens. This work underscores the importance of engineering cellular states, not just genetic constructs, to unlock genome-scale screening in non-model organisms.

## MATERIALS AND METHODS

### Strains and Culturing Conditions

*E. coli* NEB DH10β was used for cloning and cultured in LB media at 37°C. All *Candida* species strains were routinely cultured in YPD media (10 g/L yeast extract, 20 g/L peptone, 20 g/L glucose) at 30°C. YNB media was prepared using 6.7 g/L yeast nitrogen base without amino acids (Sigma-Aldrich Y0626), 5 g/L glucose, and 0.36 g/L potassium acetate unless otherwise stated. *C. albicans* SC5314 (ATCC MYA-2876) was used for most experiments in this work. *Candida* species strains including *C. albicans* Y-12983, *C. glabrata* Y-65, *C. krusei* Y-96, *C. orthopsilosis* Y-48468, *C. parapsilosis* Y-12969, and *C. dubliniensis* Y-17841 were provided by the USDA-ARS Culture Collection (NRRL). All *C. auris* isolates (AR-0381, AR-0383, AR-0385, AR-0387, and AR-1097) were provided by the CDC AR Isolate Bank.

### Plasmid Construction

All plasmids and primers are listed in Table S1 and Table S2 respectively. Cas9 containing plasmids used for transformation testing with all *Candida* species were obtained from Addgene. All gRNA seed sequences were designed in CASPER^47,48^. CRISPR-GRIT gRNAs were designed using a custom python script. All gRNA seed sequences and full CRISPR-GRIT sequences are listed in Table S3. CRISPR-GRIT plasmids harboring the *ADE2* gRNA and gRNAs used for the library screen were constructed using Golden Gate Assembly, as described in Cotter and Trinh^16^. Before transformation, 5-10 µg of pV1093-CatRNA based plasmids were linearized using KpnI and SacI in an overnight restriction digest at 37°C. The digests were purified using the Omega Biotek E.Z.N.A. Cycle Pure Kit and the total purified products were used for subsequent transformations.

### Transformation of Candida species

The protocol developed by Walther and Wendland was used as the standard LiOAc transformation method and baseline for optimization for all *Candida* species^19^. Briefly, a single, freshly streaked colony was used to inoculate 2-10 mL of YPD and grown overnight at 30°C, 250 rpm. The overnight culture was diluted in 10-50 mL of YPD to an optical density at 600nm (OD600) of 0.3 and grown at 30°C, 250 rpm for 4 hours to an OD600 of approximately 1.8. The cells were then collected by centrifugation at 900 x g for 5 minutes, washed with one-half of the culture volume of sterile water, and resuspended to an OD600 of 30 in 100 mM LiOAc dissolved in TE buffer at pH 8 (LiOAc TE). One hundred microliters of cells were aliquoted in sterile 1.5 mL centrifuge tubes, and 10 µL of 10 mg/mL herring sperm DNA (Invitrogen), 0.1-5 µg of linearized DNA, and 600 µL of 50% PEG 3350 dissolved in 100 mM LiOAc TE was added to each tube. Each tube was briefly vortexed and incubated stationary at 30°C for 20 hours. After incubating for 20 hours, tubes were vortexed to mix and placed in a 44°C water bath for 15 minutes. The tubes were then centrifuged at 900 x g for 5 minutes. The supernatant was removed carefully, and the cells were washed with 1 mL of YPD. The cells were then recovered in 2 mL of YPD at 30°C, 250 rpm for 4 hours. Following the recovery, cells were pelleted at 900 x g for 5 minutes, resuspended in sterile water and serially diluted before plating on YPD agar plates supplemented with nourseothricin (NAT). The FACT method follows this protocol exactly apart from preculturing cells in YNB supplemented with 0.36 g/L KOAc and using 100 mM LiOAc dissolved in water and titrated to pH 5 with glacial acetic acid (LiOAc pH 5). 250 µg/mL NAT was used for selection of *C. albicans* transformants. 200 µg/mL NAT was used for *C. parapsilosis, C*. *orthopsilosis* and all *C. auris* isolates. 100 µg/mL NAT was used for selection of *C. dubliniensis, C. glabrata,* and *C. krusei.* Plates were incubated at 30°C for 2-3 days before counting colonies.

### Post-transformation Spotting and 96-well Microplate Kinetic Growth Assays

The transformation procedure above was followed exactly up through washing the cells with YPD following heat shock. After washing, the cells were resuspended in 1 mL of YPD. For spotting assays, 10x serial dilutions were performed in YPD and 3 µL of each dilution was spotted on YPD agar plates. Plates were incubated for 24 hours at 30°C prior to taking images. For microplate kinetic growth assays, 20 µL of cells were added to 180 µL of YPD for a final volume of 200 µL in a 96-well microplate at an initial OD600 of ∼0.3. The microplate was incubated in an EPOCH2 plate reader at 30°C using the fastest orbital shaking setting for 24 hours, measuring the OD600 every 15 minutes.

### GSH/GSSG Assay

GSH/GSSG ratios were measured using the Promega GSH/GSSG-Glo Assay following the manufacturer’s protocol for suspension cells. Briefly, the transformation procedure above was performed up through the YPD wash step following heat shock. Cells were then resuspended in 1 mL of YPD. For untreated controls, cells were grown to an OD600 of 1.8 in their respective media and directly used for GSH/GSSG measurements. A volume of 25 µL of cells were added to a white-walled, clear-bottom 96-well plate. Subsequently, 25 µL of Total Glutathione Lysis Reagent or Oxidized Glutathione Lysis Reagent was added to the desired wells. The plate was shaken for 5 minutes on a plate shaker. Then, 50 µL of Luciferin Generation Reagent was added to each well, briefly mixed by shaking, and incubated at room temperature for 30 minutes. Finally, 100 µL of Luciferin Detection Reagent was added to each well and incubated at room temperature for 30 minutes. Luminescence was measured using a Biotek Synergy HT plate reader.

### Hydrogen Peroxide Preconditioning and Oxidative Stress Protection Assays

H_2_O_2_ was added to cultures at a final concentration of 0.4 mM during the preculture step and incubated for exactly one hour until the cells reached an OD600 of approximately 1.8. For YPD cultures, H_2_O_2_ was added when cells were at an OD600 of ∼0.9. For YNB cultures, H_2_O_2_ was added when cells were at an OD600 of ∼1.1. Following preconditioning, an aliquot of cells was washed with 1 mL of sterile PBS and resuspended to an OD600 of 1 in sterile PBS. 10x serial dilutions were performed in PBS and 3 µL of each dilution was spotted on YPD agar plates supplemented with varying concentrations of H_2_O_2_ to validate oxidative stress priming. Plates were incubated for 24 hours at 30°C prior to taking images. The remaining cells were transformed using the standard and FACT methods. Spotting viability and recovery growth kinetic assays were performed as described above.

## LC-MS/MS-based Proteomics

### Sample preparation

*C. albicans* SC5314 cells were cultured to log phase (OD600 ∼1.8) in YPD or YNB and either sampled directly (untreated samples) or sampled after transformation using the standard or FACT method immediately prior to heat shock (treated samples). Cell samples were harvested, washed, and fractionated to enrich cytoplasmic and membrane-associated proteins prior to proteomic analysis. To obtain the fractions, previously washed and snap frozen cell pellets were disrupted by bead beating using 0.5 mm zirconium oxide beads in 100 mM Tris-HCl, pH 8.0, with a Geno/Grinder (SPEX SamplePrep). Following cell disruption, samples were centrifuged at 21,000 x g to separate soluble and insoluble material. The supernatant (enriched in cytoplasmic proteins) was transferred to a fresh tube. The pelleted cellular debris (enriched in membrane-associated proteins) and beads were washed three times with 100 mM Tris-HCl to remove residual cytoplasmic proteins. Both cytoplasmic and membrane-enriched fractions were then processed in parallel using identical SDS-based protein aggregation capture (PAC) workflows, as described previously^49^. Briefly, samples were adjusted to 4% SDS and 10 mM dithiothreitol, heated to 90°C to denature proteins, clarified by centrifugation, and alkylated with 30 mM iodoacetamide. Proteins were extracted by PAC and digested *in situ* with sequencing-grade trypsin at a 1:75 (w/w) trypsin-to-protein ratio overnight, followed by a second 4 h digestion the next day. Tryptic digests were acidified to 0.5% formic acid, filtered through 10 kDa MWCO spin filters, and peptide concentrations determined by NanoDrop A205 measurements.

### LC-MS/MS Analysis

Three micrograms of tryptic peptides from each cytoplasmic and membrane fraction were analyzed by automated 1D LC-MS/MS using a Vanquish UHPLC coupled directly to an Orbitrap Q Exactive Plus mass spectrometer (Thermo Scientific). Peptides were loaded onto a 15 cm, in-house pulled nanospray column/emitter (1.7 micron Kinetex C18; Phenomenex) and separated by 180 min organic gradient, as previously described^50^ but without the use of a trapping column. Eluting peptides were measured and sequenced using data-dependent acquisition.

### MS/MS Data Analysis

Raw MS/MS spectra were analyzed using Proteome Discoverer (v3, Thermo Scientific) with the SEQUEST HT search algorithm against a *Candida albicans* proteome database along with common contaminant proteins. Peptide spectrum matches (PSMs) were required to be fully tryptic, allowing up to two missed cleavages, with carbamidomethylation of cysteine (57.0214 Da) as a static modification and oxidation of methionine (15.9949 Da) as a dynamic modification. PSMs were scored and filtered using Percolator, with false discovery rates controlled at <1% at the PSM and peptide levels. Peptide abundances were quantified by chromatographic area-under-the-curve, mapped to their corresponding proteins, and summed to estimate protein-level abundance. Missing protein abundances were imputed and all values were log_2_ transformed and normalized using variance stabilizing normalization (VSN) as implemented in the DEP R package v1.31.2 ^51^. Pearson correlations were performed using the VSN-normalized values. Differentially expressed proteins were identified using the limma R package^52^. Moderated t-statistics were calculated using empirical Bayes shrinkage and p-values were adjusted for multiple testing using the Benjamini-Hochberg false discovery rate (FDR) method. Functional enrichment analyses of protein sets were performed using DAVID^53,54^. Proteome allocation analyses were performed as described previously^55–57^.

All raw mass spectra in this study have been deposited in the MassIVE and ProteomeXchange data repositories under accession numbers MSV000100360 (MassIVE) and PXD072526 (ProteomeXchange), with data files available at ftp://MSV000100360@massive-ftp.ucsd.edu.

### Pooled CRISPR Library Screen

200 ng of individual CaVip plasmids were pooled in a total volume of 100 µL then diluted to a concentration of 10 ng/µL. 1 µL of the pooled plasmid mixture was added to a 50 µL PCR with primers CC-82 and CC-83. The amplified product was purified using Omega Biotek NGS Magbeads with a 0.8x bead:reaction ratio. 3.5 µg of the purified CaVip amplicon pool was transformed into *C. albicans* SC5314 using both the standard transformation method described above and the FACT method (preculture in YNB+KOAc and incubation in 100 mM LiOAc/PEG 3350 at pH 5). After the 4-hour recovery in 2 mL of YPD, 1 mL of cells was collected as the T0 control sample, while the remaining 1 mL was transferred to 50 mL of YPD broth containing 250 µg/mL NAT in a 500 mL baffled flask and incubated for 40 hours at 30°C, 250 rpm for gRNA depletion. The outgrowth cultures were sampled at 24 and 40 hours and pellets were stored at -20°C before genomic DNA extraction. Genomic DNA was extracted from all samples using the Thermo Scientific Yeast DNA Extraction Kit following the manufacturer’s instructions. The gRNA locus was amplified from each sample by adding 100 ng of genomic DNA in a 50 µL PCR using primers CC-674/CC-675. PCR products were purified using Omega Biotek NGS Magbeads sent for Illumina sequencing using the Amplicon-EZ service through Azenta GENEWIZ.

### Sequencing Analysis

Forward and reverse reads were paired by aligning ends in pRESTO with the AssemblePairs function. A custom Python script was then used to filter paired reads, retaining only those that contained at least the 3’ end of the tRNA (CTTATCTCGTCCA) or the 5’ end of the gRNA scaffold (GTTTTAGAGC). Individual gRNAs were identified and counted from the filtered reads.

### Statistical Analysis

Statistical analyses were performed using GraphPad Prism 10 and R. Statistical significance was estimated using two-tailed t tests for paired variables and two-way ANOVA with either Tukey’s test or Šidák corrections for multiple comparisons. Sample numbers, p-values, and the statistical test used for each analysis can be found in the respective figure legends.

## ACKNOWLEDGEMENTS

This research is financially supported in part by the ORII Seed Fund at The University of Tennessee, Knoxville, and the DOE BER Genomic Science Program (DE-SC0019412). The views, opinions, and/or findings contained in this article are those of the authors and should not be interpreted as representing the official views or policies, either expressed or implied, of the funding agencies. The mention of trade names or commercial products in this publication is solely for the purpose of providing specific information and does not imply a recommendation or endorsement by the funding agencies.

## AUTHOR CONTRIBUTIONS

Conceptualization: CT; Investigation: CC, CT; Methodology: CC, DLC, RJG, CT; Formal Analysis: CC, CT; Visualization: CC, CT; Funding Acquisition: CT; Project Administration: CT; Supervision: CT; Writing-Review & Editing: CC, RJG, CT.

