## Supplementary Figure S1-S6 and Table S1-S3 for "Cellular Stress Tolerance Governs Genetic Transformability in Recalcitrant *Candida* Species"

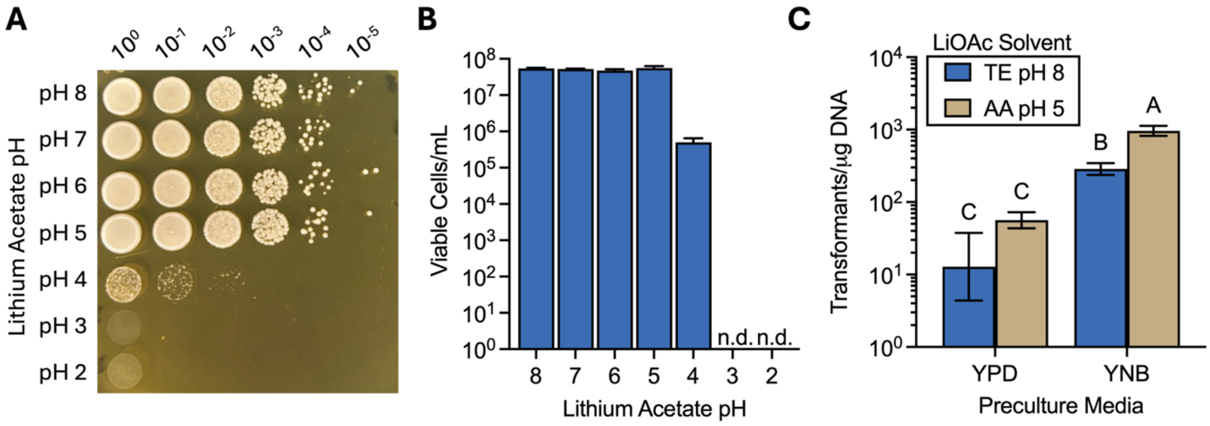


**Supplementary Figure S1. Effect of lithium acetate pH optimization on viability and transformation efficiency in *C. albicans*. (A)** Spotting assays were performed with *C. albicans* cells cultured in YNB following treatment with lithium acetate (LiOAc) titrated to varying pHs, PEG, and heat shock. Images were taken after 24 hours at 30°C and are representative of three biological replicates. **(B)** Viable cell counts were obtained by counting colony-forming units on the spotting assay plates. **(C)** Transformation efficiency of *C. albicans* precultured in either YPD or YNB and transformed using LiOAc dissolved in either TE buffer at pH 8 (TE pH 8) or in water titrated to pH 5 with acetic acid (AA pH 5). Data are the mean ± standard deviation of three biological replicates. No colonies were detected (n.d.) at pH 2 or pH 3.

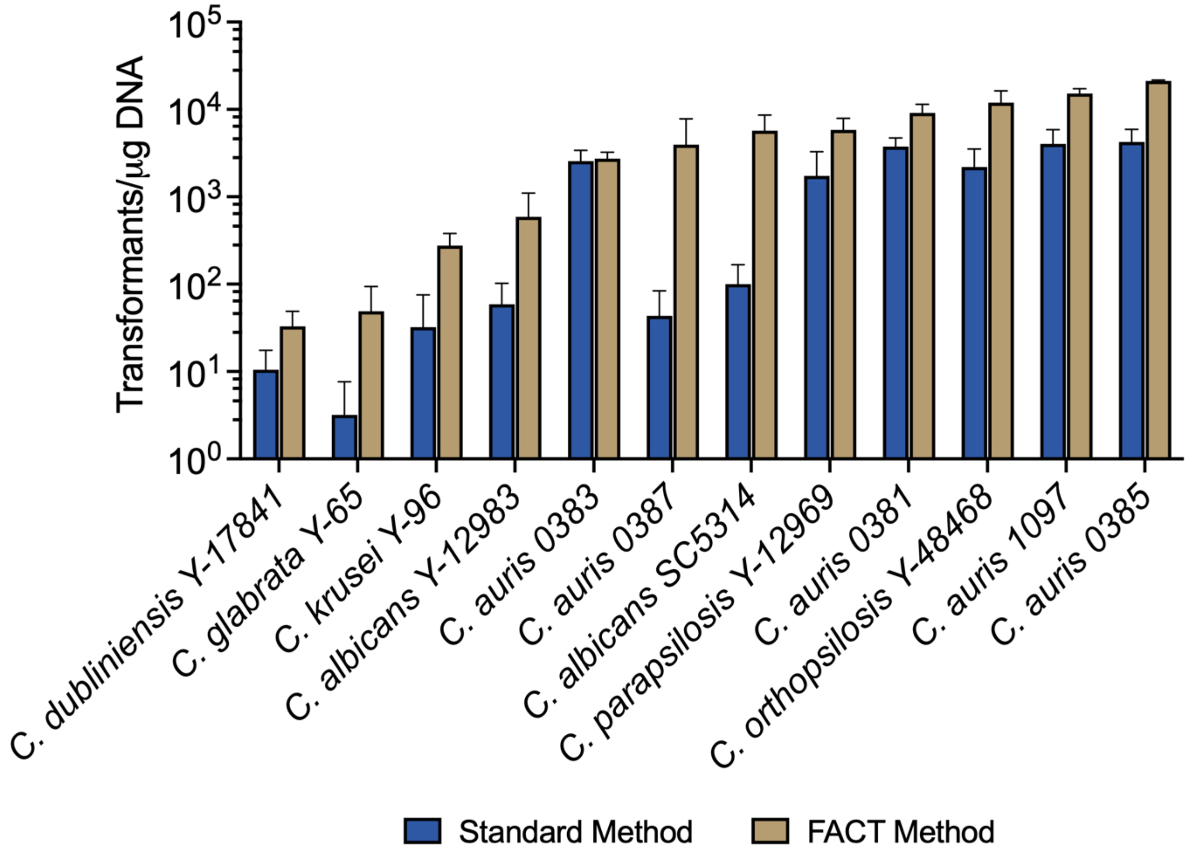


**Supplementary Figure S2. Comparison of transformation efficiencies using the standard and FACT methods among different *Candida* species.** All strains were transformed using non-targeting CRISPR-Cas plasmids or integrative DNA fragments. Data are the mean ± standard deviation of at least three biological replicates.

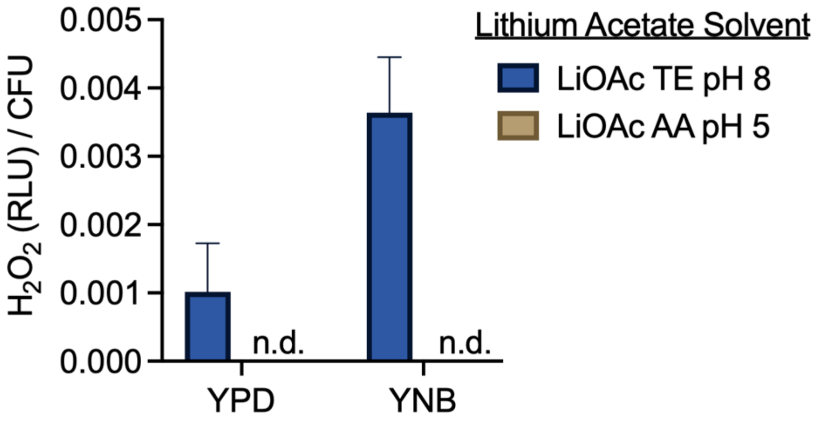


**Supplementary Figure S3. Effect of preculture media and lithium acetate solvents on the level of $H_2O_2$ detected in the transformation mixture.** $H_2O_2$ was not detected (n.d.) in samples treated with lithium acetate at pH 5 (LiOAc AA pH 5). Relative luminescence units (RLU) were normalized by viable cells per treatment (CFU) as determined by spotting assays. Data are the mean ± standard deviation of three biological replicates.

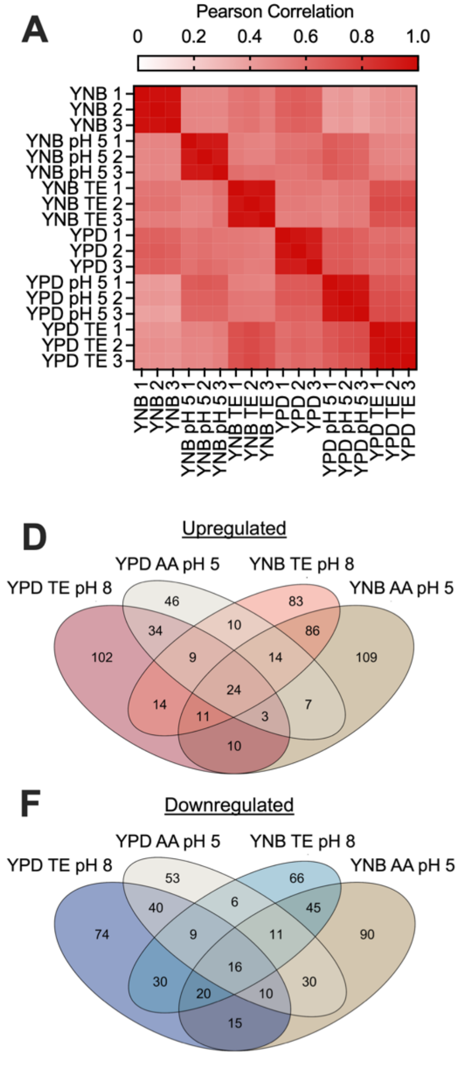


**Supplementary Figure S4. Additional Proteomics QC Metrics and Functional Profiling of Differentially Expressed Proteins. (A)** Pearson correlation across proteomics sample replicates. **(B)** PCA of YPD-precultured samples only. **(C)** PCA of YNB-precultured samples only. **(D, F)** Venn diagrams of upregulated **(D)** and downregulated **(F)** proteins across conditions relative to the respective untreated controls. **(E, G)** Heat maps presenting z-scores of summed $\log_2$ fold change values for upregulated **(E)** and **(G)** downregulated proteins belonging to each KOG functional category.

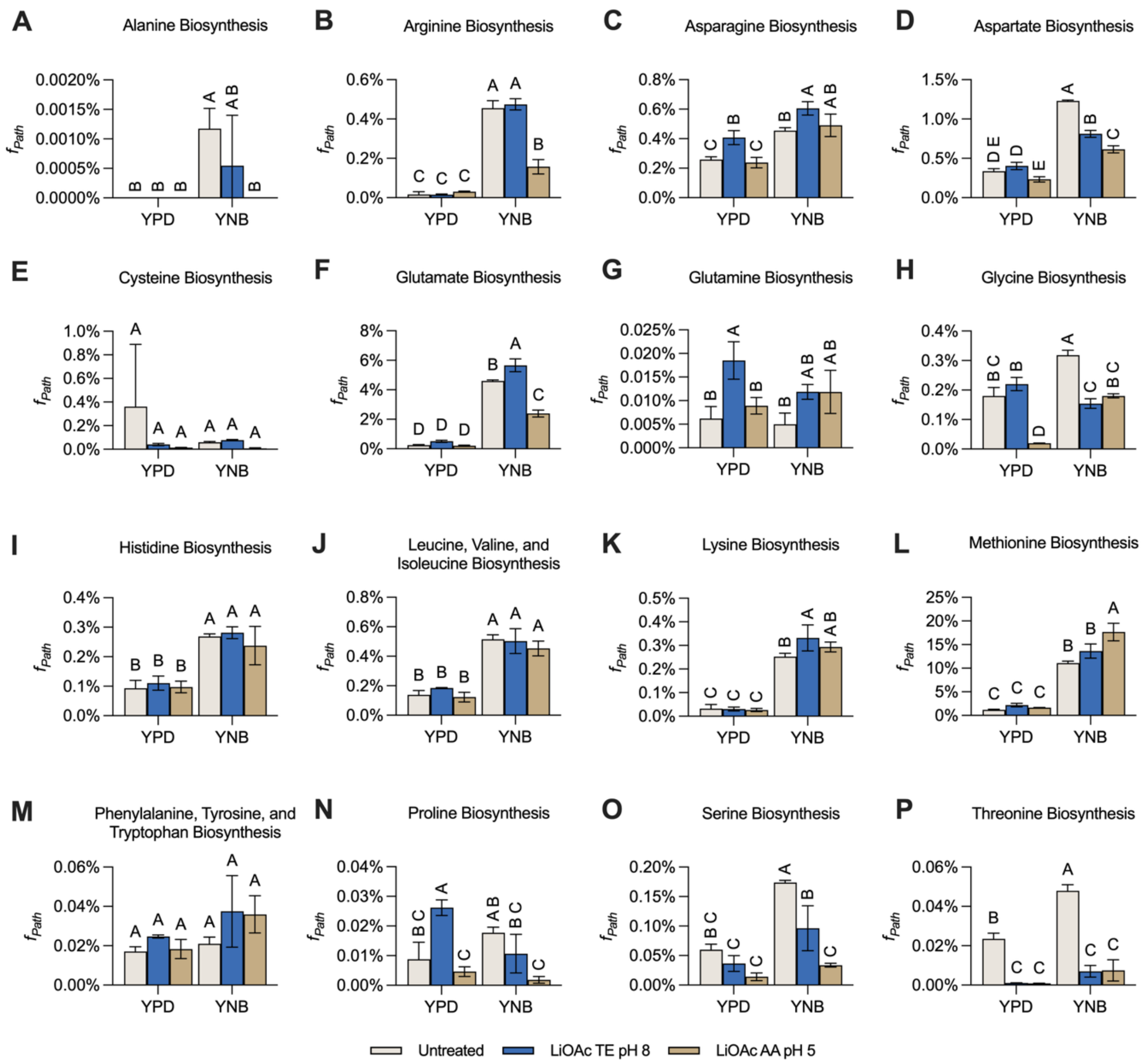


**Supplementary Figure S5. Proteome allocation towards amino acid biosynthesis before and after transformation treatment.** *C. albicans* cells were precultured in either YPD or YNB and treated with LiOAc/PEG solutions in either TE buffer at pH 8 or in water titrated to pH 5 with acetic acid prior to proteomic sample preparation. Amino acid biosynthetic pathways were curated using the Candida Genome Database[1,2]. Statistical significance was assessed using two-way ANOVA and represented with a compact letter display: groups sharing a letter are not significantly different, whereas groups with different letters are. Data are the mean ± standard deviation of three biological replicates.

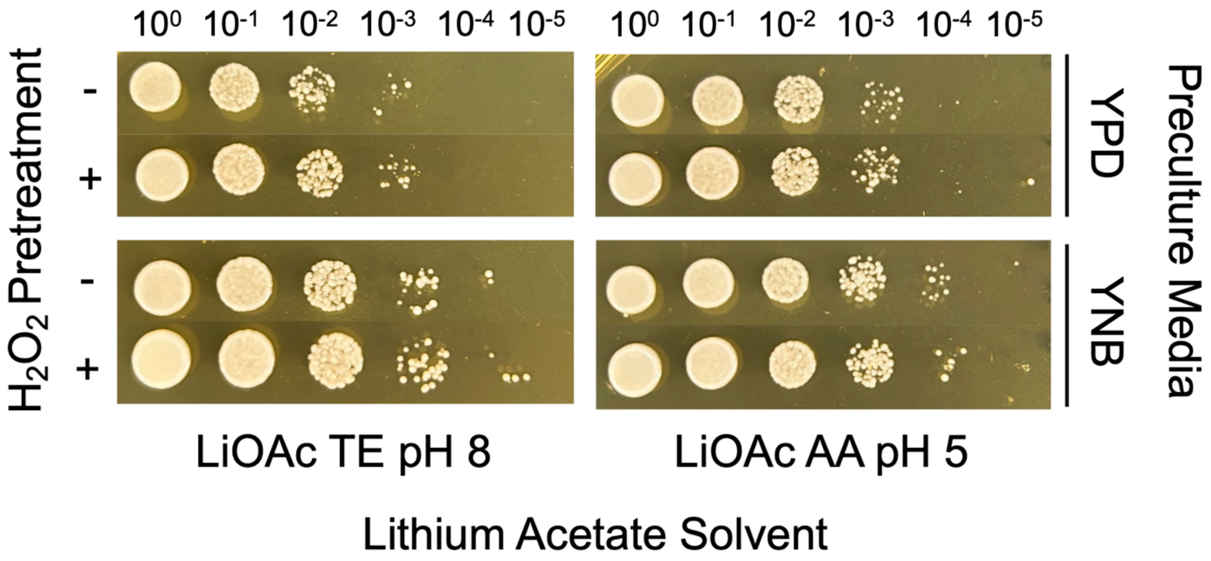


**Supplementary Figure S6. Post-transformation viability following $H_2O_2$ pretreatment.** Cells were treated with or without 0.4 mM $H_2O_2$ during the preculture step, one hour prior to cell collection. Cells were then transformed using lithium acetate dissolved in indicated solvents and following heat shock, cells were serially diluted and spotted on YPD plates to evaluate viability. No differences in post-transformation viability were observed between $H_2O_2$ treated and untreated samples. Images were taken after 24 hours at 30°C and are representative of three biological replicates.

**Supplementary Table S1:** List of plasmids used in this study.

| Plasmid | Description | Source |
|---|---|---|
| pV1093-CatRNA | pV1093 based integrative CRISPR-Cas9 system with a tRNA-Ala at the 3' end of the SNR52 promoter for increased editing efficiency | Cotter & Trinh[1] |
| CRISPR-GRIT ADE2 | pV1093 based integrative CRISPR-Cas9 system with a CRISPR-GRIT ADE2 targeting gRNA. Also referred to as CaVip-ADE2. | Cotter & Trinh[1] |
| pCT-tRNA | ctrc | Lombardi et al. (Addgene #133813)[2] |
| pCP-tRNA | CRISPR-Cas9 plasmid for genome editing *Candida parapsilosis*. Used for transformation testing of *C. parapsilosis, C. orthopsilosis,* and *C. krusei*. | Lombardi et al. (Addgene #133812)[2] |
| pCE27 | Part 1 of a split marker CRISPR-Cas9 system containing Cas9 for genome editing *Candida auris* | Ennis et al. (Addgene #174405)[3] |
| pCE35 | Part 2 of a split marker CRISPR-Cas9 system containing the sgRNA for genome editing *Candida auris* | Ennis et al. (Addgene #174409)[3] |
| CaVip-URA3 | pV1093-CatRNA based integrative CRISPR-Cas9 system with a CRISPR-GRIT gRNA targeting URA3 | Cotter & Trinh[1] |
| CaVip-1 | pV1093-CatRNA based integrative CRISPR-Cas9 system with a CRISPR-GRIT gRNA targeting RPS15 (Guide #1) | This work |
| CaVip-2 | pV1093-CatRNA based integrative CRISPR-Cas9 system with a CRISPR-GRIT gRNA targeting RPS15 (Guide #2) | This work |
| CaVip-3 | pV1093-CatRNA based integrative CRISPR-Cas9 system with a CRISPR-GRIT gRNA targeting RPS15 (Guide #3) | This work |
| CaVip-4 | pV1093-CatRNA based integrative CRISPR-Cas9 system with a CRISPR-GRIT gRNA targeting RPS15 (Guide #4) | This work |
| CaVip-5 | pV1093-CatRNA based integrative CRISPR-Cas9 system with a CRISPR-GRIT gRNA targeting TRL1 (Guide #1) | This work |
| CaVip-6 | pV1093-CatRNA based integrative CRISPR-Cas9 system with a CRISPR-GRIT gRNA targeting TRL1 (Guide #2) | This work |
| CaVip-7 | pV1093-CatRNA based integrative CRISPR-Cas9 system with a CRISPR-GRIT gRNA targeting TRL1 (Guide #3) | This work |

| | | |
|---|---|---|
| CaVip-8 | pV1093-CatRNA based integrative CRISPR-Cas9 system with a CRISPR-GRIT gRNA targeting TRL1 (Guide #4) | This work |
| CaVip-9 | pV1093-CatRNA based integrative CRISPR-Cas9 system with a CRISPR-GRIT gRNA targeting RPB2 (Guide #1) | This work |
| CaVip-10 | pV1093-CatRNA based integrative CRISPR-Cas9 system with a CRISPR-GRIT gRNA targeting RPB2 (Guide #2) | This work |
| CaVip-11 | pV1093-CatRNA based integrative CRISPR-Cas9 system with a CRISPR-GRIT gRNA targeting RPB2 (Guide #3) | This work |
| CaVip-12 | pV1093-CatRNA based integrative CRISPR-Cas9 system with a CRISPR-GRIT gRNA targeting RPB2 (Guide #4) | This work |
| CaVip-13 | pV1093-CatRNA based integrative CRISPR-Cas9 system with a CRISPR-GRIT gRNA targeting RAD52 (Guide #1) | This work |
| CaVip-14 | pV1093-CatRNA based integrative CRISPR-Cas9 system with a CRISPR-GRIT gRNA targeting RAD52 (Guide #2) | This work |
| CaVip-15 | pV1093-CatRNA based integrative CRISPR-Cas9 system with a CRISPR-GRIT gRNA targeting RAD52 (Guide #3) | This work |
| CaVip-16 | pV1093-CatRNA based integrative CRISPR-Cas9 system with a CRISPR-GRIT gRNA targeting RAD52 (Guide #4) | This work |
| CaVip-17 | pV1093-CatRNA based integrative CRISPR-Cas9 system with a CRISPR-GRIT gRNA targeting RAD51 (Guide #1) | This work |
| CaVip-18 | pV1093-CatRNA based integrative CRISPR-Cas9 system with a CRISPR-GRIT gRNA targeting RAD51 (Guide #2) | This work |
| CaVip-19 | pV1093-CatRNA based integrative CRISPR-Cas9 system with a CRISPR-GRIT gRNA targeting RAD51 (Guide #3) | This work |
| CaVip-20 | pV1093-CatRNA based integrative CRISPR-Cas9 system with a CRISPR-GRIT gRNA targeting RAD51 (Guide #4) | This work |
| CaVip-21 | pV1093-CatRNA based integrative CRISPR-Cas9 system with a CRISPR-GRIT gRNA targeting LIG4 (Guide #1) | This work |
| CaVip-22 | pV1093-CatRNA based integrative CRISPR-Cas9 system with a CRISPR-GRIT gRNA targeting LIG4 (Guide #2) | This work |
| CaVip-23 | pV1093-CatRNA based integrative CRISPR-Cas9 system with a CRISPR-GRIT gRNA targeting LIG4 (Guide #3) | This work |

| | | |
|---|---|---|
| CaVip-24 | pV1093-CatRNA based integrative CRISPR-Cas9 system with a CRISPR-GRIT gRNA targeting LIG4 (Guide #4) | This work |
| CaVip-25 | pV1093-CatRNA based integrative CRISPR-Cas9 system with a CRISPR-GRIT gRNA targeting ALS1 (Guide #1) | This work |
| CaVip-26 | pV1093-CatRNA based integrative CRISPR-Cas9 system with a CRISPR-GRIT gRNA targeting ALS1 (Guide #2) | This work |
| CaVip-27 | pV1093-CatRNA based integrative CRISPR-Cas9 system with a CRISPR-GRIT gRNA targeting FLO8 (Guide #1) | This work |
| CaVip-28 | pV1093-CatRNA based integrative CRISPR-Cas9 system with a CRISPR-GRIT gRNA targeting FLO8 (Guide #2) | This work |
| CaVip-29 | pV1093-CatRNA based integrative CRISPR-Cas9 system with a CRISPR-GRIT gRNA targeting ACE2 (Guide #1) | This work |
| CaVip-30 | pV1093-CatRNA based integrative CRISPR-Cas9 system with a CRISPR-GRIT gRNA targeting ACE2 (Guide #2) | This work |
| CaVip-Null-1 | pV1093-CatRNA based integrative CRISPR-Cas9 system with a null targeting guide (Guide #1) | This work |
| CaVip-Null-2 | pV1093-CatRNA based integrative CRISPR-Cas9 system with a null targeting guide (Guide #2) | This work |
| CaVip-Null-3 | pV1093-CatRNA based integrative CRISPR-Cas9 system with a null targeting guide (Guide #3) | This work |

**Supplementary Table S2:** List of primers used in this study.

| Primer Name | Sequence (5'-3') | Description |
|---|---|---|
| CC-82 | GCCACTACTACCACTGGGAG | Forward primer for PCR linearization of pV1093-CatRNA based plasmids for library. |
| CC-83 | CGAACACTCTTGTGCTGATC | Reverse primer for PCR linearization of pV1093-CatRNA based plasmids for library. |
| CC-674 | GAGTTCGACTCTTATCTCGTCC | Forward primer for amplifying CRISPR-GRIT gRNA locus from genomic DNA for library sequencing. |
| CC-675 | CAAGTTGATAACGGACTAGCC | Reverse primer for amplifying CRISPR-GRIT gRNA locus from genomic DNA for library sequencing. |
| CC-534 | GACGGCACGGCCACGCGTTTAAACCGCC | Universal A forward primer for amplifying Cas9 fragment from pCE35 for *C. auris* split marker system transformation[3]. |
| CC-535 | CCCGCCAGGCGCTGGGGTTTAAACACCG | Universal B reverse primer for amplifying Cas9 fragment from pCE35 for *C. auris* split marker system transformation[3]. |
| CC-537 | AGGTGATGCTGAAGCTATTGAAG | Universal C forward primer for amplifying gRNA fragment from pCE27 for *C. auris* split marker system transformation[3]. |
| CC-538 | TTATTTCTGCAAAAGCTTCTTTAC | Universal C reverse primer for amplifying gRNA fragment from pCE27 for *C. auris* split marker system transformation[3]. |
| CaVip-1-GRIT_FW | AAACGTCTCTGTCCATGGTTGGTCACTACTTGGGTGAATTCTCCATTACCTATACTCCAGTTAGAtaataaAGCTGGTAATGCTTCTTC | Forward primer for construction of CaVip-1 GRIT gRNA |
| CaVip-1-GRIT_RV | TTTCGTCTCCAAAACTGTCTAACTGGAGTATAGGTCACATCAAGTTTATCTCAATGGCATGAATTTAGAAGAAGCATTACCAGCTttat | Reverse primer for construction of CaVip-1 GRIT gRNA |

| | | |
|---|---|---|
| CaVip-2-GRIT_FW | AAACGTCTCTGTCCATCAAAAAATTGAGAGCTGCCAGAGCTGCCACTGAACCAAATGAAAGACCAtaataaCAAAACTCACTTGAGAAA | Forward primer for construction of CaVip-2 GRIT gRNA |
| CaVip-2-GRIT_RV | TTTCGTCTCCAAAACTTGTCAAAACTCACTTGAGACAGAACCAATCATTTCTGGAACAACAATCATGTTTCTCAAGTGAGTTTTGttat | Reverse primer for construction of CaVip-2 GRIT gRNA |
| CaVip-3-GRIT_FW | AAACGTCTCTGTCCAGAAACATGATTGTTGTTCCAGAAATGATTGGTTCTGTTGTTGGTGTCTACtaataaAGTTTTCAACACTGTTGA | Forward primer for construction of CaVip-3 GRIT gRNA |
| CaVip-3-GRIT_RV | TTTCGTCTCCAAAACTTGTAGACACCAACAACAGACCAAGTAGTGACCAACCATTTCTGGTTTAATTTCAACAGTGTTGAAAACTttat | Reverse primer for construction of CaVip-3 GRIT gRNA |
| CaVip-4-GRIT_FW | AAACGTCTCTGTCCAGTAAAGTTTTCAACACTGTTGAAATTAAACCAGAAATGGTTGGTCACTACtaataaATTCTCCATTACCTATAC | Forward primer for construction of CaVip-4 GRIT gRNA |
| CaVip-4-GRIT_RV | TTTCGTCTCCAAAACAAGTAGTGACCAACCATTTCAAGCATTACCAGCTCTACCGTGTCTAACTGGAGTATAGGTAATGGAGAATttat | Reverse primer for construction of CaVip-4 GRIT gRNA |
| CaVip-5-GRIT_FW | AAACGTCTCTGTCCAAAAGTGAAATTGAACCAATTAATAAACAACTCCATATTACCATTGGTTGTtaataaAGCTACTGCTGTTGAGTC | Forward primer for construction of CaVip-5 GRIT gRNA |
| CaVip-5-GRIT_RV | TTTCGTCTCCAAAACGCCAGCTACTGCTGTTGAGTGATTGTCATACAATTCTTCCAATGTTATGTTTGACTCAACAGCAGTAGCTttat | Reverse primer for construction of CaVip-5 GRIT gRNA |
| CaVip-6-GRIT_FW | AAACGTCTCTGTCCAATGACGTTCTACAAAGTGAAATTGAACCAATTAATAAACAACTCCATATTtaataaTTGTATCCCGCCAGCTAC | Forward primer for construction of CaVip-6 GRIT gRNA |

| | | |
|---|---|---|
| CaVip-6-GRIT_RV | TTTCGTCTCCAAAACTTGGTTGTATCCCGCCAGCTATTCTTCCAATGTTATGTTTGACTCAACAGCAGTAGCTGGCGGGATACAAttat | Reverse primer for construction of CaVip-6 GRIT gRNA |
| CaVip-7-GRIT_FW | AAACGTCTCTGTCCACTCAGATCAAATTAGATTCTTGGAAATTTTTGGAATGGGATTATGGCAAAtaataaTCAACTTCCAATTCAAGC | Forward primer for construction of CaVip-7 GRIT gRNA |
| CaVip-7-GRIT_RV | TTTCGTCTCCAAAACAAGGTTTGCCATAATCCCATTAGTATCATTATTTAATGTAAACAATCCACGCGCTTGAATTGGAAGTTGAttat | Reverse primer for construction of CaVip-7 GRIT gRNA |
| CaVip-8-GRIT_FW | AAACGTCTCTGTCCAATGGTCGAGAAATTGAAGGGTTTGTGATAAGATGTCACCGCCAACTGCATtaataaTGACACTGACGGGGATTG | Forward primer for construction of CaVip-8 GRIT gRNA |
| CaVip-8-GRIT_RV | TTTCGTCTCCAAAACATGGTGACACTGACGGGGATATGGCTGTTCAAATTTATACTTGAAAAAAAAGCAATCCCCGTCAGTGTCAttat | Reverse primer for construction of CaVip-8 GRIT gRNA |
| CaVip-9-GRIT_FW | AAACGTCTCTGTCCAGAATTAAGCCAACCACCAGTGGCAATACTCATACATACACACATTGTGAAtaataaGTCTATGATATTGGGTGT | Forward primer for construction of CaVip-9 GRIT gRNA |
| CaVip-9-GRIT_RV | TTTCGTCTCCAAAACTCCGTCTATGATATTGGGTGTATGATCTGGGAACGGAATAATTGAGGCGGCAACACCCAATATCATAGACttat | Reverse primer for construction of CaVip-9 GRIT gRNA |
| CaVip-10-GRIT_FW | AAACGTCTCTGTCCAATGGTGGAGACTTTAATGTTACTTTGGCTGTTAAATCACAGACAATCACCtaataaAAGATACTCGTTGGCTAC | Forward primer for construction of CaVip-10 GRIT gRNA |
| CaVip-10-GRIT_RV | TTTCGTCTCCAAAACTCGGTGATTGTCTGTGATTTTCATGGCTTTTCTTTGCTCACCCCAATTACCAGTAGCCAACGAGTATCTTttat | Reverse primer for construction of CaVip-10 GRIT gRNA |

| | | |
|---|---|---|
| CaVip-11-GRIT_FW | AAACGTCTCTGTCCAGATTAAAACACGGTACATATGAGAAATTAGATGAAGATGGGTTGATCGCCtaataaCAGAGTTAGTGGTGAAGA | Forward primer for construction of CaVip-11 GRIT gRNA |
| CaVip-11-GRIT_RV | TTTCGTCTCCAAAACCAGGGGCGATCAACCCATCTCAGGTATAGGAGTTGTTTACCAATAATAATATCTTCACCACTAACTCTGttat | Reverse primer for construction of CaVip-11 GRIT gRNA |
| CaVip-12-GRIT_FW | AAACGTCTCTGTCCAGATTAAAACACGGTACATATGAGAAATTAGATGAAGATGGGTTGATCGCCtaataaCAGAGTTAGTGGTGAAGA | Forward primer for construction of CaVip-12 GRIT gRNA |
| CaVip-12-GRIT_RV | TTTCGTCTCCAAAACAGGGGCGATCAACCCATCTTCAGGTATAGGAGTTGTTTACCAATAATAATATCTTCACCACTAACTCTGttat | Reverse primer for construction of CaVip-12 GRIT gRNA |
| CaVip-13-GRIT_FW | AAACGTCTCTGTCCACAATTAAACGAAGTGATATTAAAAATAGTGTCGCTACGACACCGTCACCAtaataaAAACACATCTAGTAATCG | Forward primer for construction of CaVip-13 GRIT gRNA |
| CaVip-13-GRIT_RV | TTTCGTCTCCAAAACAATGGTGACGGTGTCGTAGCTATTGTTTGATGTTATCATGGGTTTATTAATTCGATTACTAGATGTGTTTttat | Reverse primer for construction of CaVip-13 GRIT gRNA |
| CaVip-14-GRIT_FW | AAACGTCTCTGTCCAAAGAACAAAAGAGACAACAAGAAGACGCGTCAAGGTTGGATAATTCAGGTtaataaGCAGCACCAGCAACAAAA | Forward primer for construction of CaVip-14 GRIT gRNA |
| CaVip-14-GRIT_RV | TTTCGTCTCCAAAACGTTGACCTGAATTATCCAACGAATTGATCCGTTTGAAACTACGTTTGAGTTATTTTGTTGCTGGTGCTGCttat | Reverse primer for construction of CaVip-14 GRIT gRNA |
| CaVip-15-GRIT_FW | AAACGTCTCTGTCCAGACCACCACAACAACAACCTCAGCAACCTCAGCAACCTCAACCCAACCAAtaataaTCCCTTTCGACCAGACGA | Forward primer for construction of CaVip-15 GRIT gRNA |

| | | |
|---|---|---|
| CaVip-15-GRIT_RV | TTTCGTCTCCAAAACCTGTTGGTTGGGTTGAGGTTCAAGTGGTTGATTCAAACGACGAGCTCGTGACTCGTCTGGTCGAAAGGGAttat | Reverse primer for construction of CaVip-15 GRIT gRNA |
| CaVip-16-GRIT_FW | AAACGTCTCTGTCCACTCCGCAACCACGACCACCACAACAACAACCTCAGCAACCTCAGCAACCTtaataaCCAACAGAGGCTTCCCTT | Forward primer for construction of CaVip-16 GRIT gRNA |
| CaVip-16-GRIT_RV | TTTCGTCTCCAAAACAACCAACAGAGGCTTCCCTTTCAAACGACGAGCTCGTGACTCGTCTGGTCGAAAGGGAAGCCTCTGTTGGttat | Reverse primer for construction of CaVip-16 GRIT gRNA |
| CaVip-17-GRIT_FW | AAACGTCTCTGTCCAAAGTTGATGGTATGTCTGGTATGTTTAATCCTGATCCTAAGAAACCAATTtaataaCATTATTGCCCATTCATC | Forward primer for construction of CaVip-17 GRIT gRNA |
| CaVip-17-GRIT_RV | TTTCGTCTCCAAAACCCAATTGGTTTCTTAGGATCCTCTTCCCTTTTTAAACGATAATCTAGTTGTTGATGAATGGGCAATAATGttat | Reverse primer for construction of CaVip-17 GRIT gRNA |
| CaVip-18-GRIT_FW | AAACGTCTCTGTCCACTCAAGTTGATGGTATGTCTGGTATGTTTAATCCTGATCCTAAGAAACCAtaataaTAACATTATTGCCCATTC | Forward primer for construction of CaVip-18 GRIT gRNA |
| CaVip-18-GRIT_RV | TTTCGTCTCCAAAACCAATTGGTTTCTTAGGATCATTCCCTTTTTAAACGATAATCTAGTTGTTGATGAATGGGCAATAATGTTAttat | Reverse primer for construction of CaVip-18 GRIT gRNA |
| CaVip-19-GRIT_FW | AAACGTCTCTGTCCATTGTTGCTCAAGTTGATGGTATGTCTGGTATGTTTAATCCTGATCCTAAGtaataaTGGGGGTAACATTATTGC | Forward primer for construction of CaVip-19 GRIT gRNA |
| CaVip-19-GRIT_RV | TTTCGTCTCCAAAACATTGGGGGTAACATTATTGCTTTTAAACGATAATCTAGTTGTTGATGAATGGGCAATAATGTTACCCCCAttat | Reverse primer for construction of CaVip-19 GRIT gRNA |

| | | |
|---|---|---|
| CaVip-20-GRIT_FW | AAACGTCTCTGTCCATTAGAACAGGTAAATCACAATTATGTCACACGTTAGCCGTTACGTGTCAGtaataaTGATATGGGTGGTGGTGA | Forward primer for construction of CaVip-20 GRIT gRNA |
| CaVip-20-GRIT_RV | TTTCGTCTCCAAAACATTGATATGGGTGGTGGTGATACCTTCAGTATCAATGTAAAGACATTTTCCTTCACCACCACCCATATCAttat | Reverse primer for construction of CaVip-20 GRIT gRNA |
| CaVip-21-GRIT_FW | AAACGTCTCTGTCCAATGACTTACAAAAGACATTACAATTTTCAACAAATCCCGATTTGAGACTAtaataaGCAGTTAGCTATACATCC | Forward primer for construction of CaVip-21 GRIT gRNA |
| CaVip-21-GRIT_RV | TTTCGTCTCCAAAACTCTGCAGTTAGCTATACATCTTTCTGACAACTGTGGTTTAAACTTGAAACAAGGATGTATAGCTAACTGCttat | Reverse primer for construction of CaVip-21 GRIT gRNA |
| CaVip-22-GRIT_FW | AAACGTCTCTGTCCAATAAAAATGGTGGGTCTTCGAGAATTCCACATTTTGTTACTGAGGCATTCtaataaTTCAATCAAGATGAATTA | Forward primer for construction of CaVip-22 GRIT gRNA |
| CaVip-22-GRIT_RV | TTTCGTCTCCAAAACGAACGAATGCCTCAGTAACAACCTAAACTTGTAATCGTCAGGATCGGGAATATAATTCATCTTGATTGAAttat | Reverse primer for construction of CaVip-22 GRIT gRNA |
| CaVip-23-GRIT_FW | AAACGTCTCTGTCCAAAAGTGAACTTGAAGACCTTCGTAATGGATTTGATTGGGGTGATCTTAAAtaataaCTACTTGTTCAAAGGTTT | Forward primer for construction of CaVip-23 GRIT gRNA |
| CaVip-23-GRIT_RV | TTTCGTCTCCAAAACAGGTTTAAGATCACCCCAATCACTCAAGTTATTCCCACACACGTAAAATGACAAACCTTTGAACAAGTAGttat | Reverse primer for construction of CaVip-23 GRIT gRNA |
| CaVip-24-GRIT_FW | AAACGTCTCTGTCCATATTGAAGAATGTACAGCTGAAGTATGAAATTGATGGGTTCAGAAACCCAtaataaAAAAGTCAAACCAGAGTA | Forward primer for construction of CaVip-24 GRIT gRNA |

| | | |
|---|---|---|
| CaVip-24-GRIT_RV | TTTCGTCTCCAAAACATCTGGGTTTCTGAACCCATCAAGATCTAAATTTTCACCAAATTTTTCTAAATACTCTGGTTTGACTTTTttat | Reverse primer for construction of CaVip-24 GRIT gRNA |
| CaVip-25-GRIT_FW | AAACGTCTCTGTCCAAATCTACTTCCACTGAAATTGAAATTGTAACAACCAGTTCTACTAAAGTTtaataaTGTCGTTTCTTCTAATAC | Forward primer for construction of CaVip-25 GRIT gRNA |
| CaVip-25-GRIT_RV | TTTCGTCTCCAAAACCCTGTCGTTTCTTCTAATACCTCTGGTATTTGTTGGTTCACTAGTCAAATCAGTATTAGAAGAAACGACAttat | Reverse primer for construction of CaVip-25 GRIT gRNA |
| CaVip-26-GRIT_FW | AAACGTCTCTGTCCAATGGTGATAAAGATATCTCAATTGATGTTGAGTTTGAAAAGTCAACCGTTtaataaTGGATATTTGTATGCTTC | Forward primer for construction of CaVip-26 GRIT gRNA |
| CaVip-26-GRIT_RV | TTTCGTCTCCAAAACAGTGGATATTTGTATGCTTCTTGTGACCTTATTGAGACTTGGCATAACTCTGGAAGCATACAAATATCCAttat | Reverse primer for construction of CaVip-26 GRIT gRNA |
| CaVip-27-GRIT_FW | AAACGTCTCTGTCCATTGGTGCAAGATCAAATCAACAGCTGACTCCACAACAAAATGCTGCACCAtaataaGCCACATCCTTCTCAACA | Forward primer for construction of CaVip-27 GRIT gRNA |
| CaVip-27-GRIT_RV | TTTCGTCTCCAAAACCTGCCACATCCTTCTCAACATCTGGAAATTATGTTGAGCTTGTGCTTGACCTTGTTGAGAAGGATGTGGCttat | Reverse primer for construction of CaVip-27 GRIT gRNA |
| CaVip-28-GRIT_FW | AAACGTCTCTGTCCACATCTGGACAAACAACACCAAACATGTCACAACCTCCTAGTGCTGGCACGtaataaGCAGCCTCAACCAACAGA | Forward primer for construction of CaVip-28 GRIT gRNA |
| CaVip-28-GRIT_RV | TTTCGTCTCCAAAACAAGCAGCCTCAACCAACAGAGCTGTTGCTTGTCTTGTAATTGACGCATTTGCTCTGTTGGTTGAGGCTGCttat | Reverse primer for construction of CaVip-28 GRIT gRNA |

| | | |
|---|---|---|
| CaVip-29-GRIT_FW | AAACGTCTCTGTCCAATTTCAACACCTTCTTACCTCCACCTACTCCGCCTAATTTGATTAATGGAtaataaTTGGAACTCATCGCCAGA | Forward primer for construction of CaVip-29 GRIT gRNA |
| CaVip-29-GRIT_RV | TTTCGTCTCCAAAACGATTGGAACTCATCGCCAGAGTTGCAATCTTCCTGGGGAAGGAGAATGTGGTTCTGGCGATGAGTTCCAAttat | Reverse primer for construction of CaVip-29 GRIT gRNA |
| CaVip-30-GRIT_FW | AAACGTCTCTGTCCATGCATTTATCACCTTTGAAAAAACAATTACCAAACACTCCCACAAAGCAAtaataaCACCATTGAATGGAGTCC | Forward primer for construction of CaVip-30 GRIT gRNA |
| CaVip-30-GRIT_RV | TTTCGTCTCCAAAACCTGTCACCATTGAATGGAGTGTAATGGTTGCTTTGAGTTTGGTGATATAACTGGACTCCATTCAATGGTGttat | Reverse primer for construction of CaVip-30 GRIT gRNA |
| CC-597 | gtccaATGCCCCTCGGGACCCCATGg | Nontargeting Guide 1 FW |
| CC-598 | aaaacCATGGGGTCCCGAGGGGCATt | Nontargeting Guide 1 RV |
| CC-599 | gtccaTTGGGATAGGGCACAGTACAg | Nontargeting Guide 2 FW |
| CC-600 | aaaacTGTACTGTGCCCTATCCCAAt | Nontargeting Guide 2 RV |
| CC-601 | gtccaGGGGACCCCGGAGAGTGCATg | Nontargeting Guide 3 FW |
| CC-602 | aaaacATGCACTCTCCGGGGTCCCCt | Nontargeting Guide 3 RV |

**Supplementary Table S3:** List of gRNAs and CRISPR-GRIT sequences used in this study.

| Plasmid | gRNA Seed Sequence (5'-3') | CRISPR-GRIT Sequence (Repair Template + Seed) |
|---|---|---|
| CaVip-ADE2 | AACACCAATGACAGGCAATG | GAATCAACCCCATCTAATGTAGATCCCTTAACTGGAACACCAATGACAGGctattaCATGGCCGCCACCATACCTGGCAAATGAGCAGCACCACCTGCACCAGCAAAACACCAATGACAGGCAATG |
| CaVip-URA3 | CCACCAACCAAGAGCCAAGA | TTGAAGGATTAAAACAGGGAGCTAAAGAAACCACCACCAACCAAGAGCCAtaataaATTGATGTTAGCTGAATTATCATCAGTGGGATCATTAGCATATGGAGAATCCACCAACCAAGAGCCAAGA |
| CaVip-1 | ACCTATACTCCAGTTAGACA | TGGTTGGTCACTACTTGGGTGAATTCTCCATTACCTATACTCCAGTTAGAtaataaAGCTGGTAATGCTTCTTCTAAATTCATGCCATTGAGATAAACTTGATGTGACCTATACTCCAGTTAGACA |
| CaVip-2 | TCTCAAGTGAGTTTTGACAA | TCAAAAAATTGAGAGCTGCCAGAGCTGCCACTGAACCAAATGAAAGACCAtaataaCAAAACTCACTTGAGAAACATGATTGTTGTTCCAGAAATGATTGGTTCTGTCTCAAGTGAGTTTTGACAA |
| CaVip-3 | TCTGTTGTTGGTGTCTACAA | GAAACATGATTGTTGTTCCAGAAATGATTGGTTCTGTTGTTGGTGTCTACtaataaAGTTTTCAACACTGTTGAAATTAAACCAGAAATGGTTGGTCACTACTTGGTCTGTTGTTGGTGTCTACAA |
| CaVip-4 | GAAATGGTTGGTCACTACTT | GTAAAGTTTTCAACACTGTTGAAATTAAACCAGAAATGGTTGGTCACTACtaataaATTCTCCATTACCTATACTCCAGTTAGACACGGTAGAGCTGGTAATGCTTGAAATGGTTGGTCACTACTT |
| CaVip-5 | ACTCAACAGCAGTAGCTGGC | AAAGTGAAATTGAACCAATTAATAAACAACTCCATATTACCATTGGTTGTtaataaAGCTACTGCTGTTGAGTCAAACATAACATTGGAAGAATTGTATGACAATCACTCAACAGCAGTAGCTGGC |
| CaVip-6 | AGCTGGCGGGATACAACCAA | ATGACGTTCTACAAAGTGAAATTGAACCAATTAATAAACAACTCCATATTtaataaTTGTATCCCGCCAGCTACTGCTGTTGAGTCAAACATAACATTGGAAGAATAGCTGGCGGGATACAACCAA |

| | | |
|---|---|---|
| CaVip-7 | ATGGGATTATGGCA AACCTT | CTCAGATCAAATTAGATTCTTGGAAATTTTTGGAATGGGATTATGGCAAAta ataaTCAACTTCCAATTCAAGCGCGTGGATTGTTTACATTAAATAATGATACT AATGGGATTATGGCAAACCTT |
| CaVip-8 | ATCCCCGTCAGTGT CACCAT | ATGGTCGAGAAATTGAAGGGTTTGTGATAAGATGTCACCGCCAACTGCATta ataaTGACACTGACGGGGATTGCTTTTTTTTCAAGTATAAATTTGAACAGCCA TATCCCCGTCAGTGTCACCAT |
| CaVip-9 | CACCCAATATCATA GACGGA | GAATTAAGCCAACCACCAGTGGCAATACTCATACATACACACATTGTGAAta ataaGTCTATGATATTGGGTGTTGCCGCCTCAATTATTCCGTTCCCAGATCATA CACCCAATATCATAGACGGA |
| CaVip-10 | AAATCACAGACAAT CACCGA | ATGGTGGAGACTTTAATGTTACTTTGGCTGTTAAATCACAGACAATCACCtaa taaAAGATACTCGTTGGCTACTGGTAATTGGGGTGAGCAAAGAAAAGCCATG AAAATCACAGACAATCACCGA |
| CaVip-11 | AGATGGGTTGATCG CCCCTG | GATTAAAACACGGTACATATGAGAAATTAGATGAAGATGGGTTGATCGCCt aataaCAGAGTTAGTGGTGAAGATATTATTATTGGTAAAACAACTCCTATACC TGAGATGGGTTGATCGCCCCTG |
| CaVip-12 | AAGATGGGTTGATC GCCCCT | GATTAAAACACGGTACATATGAGAAATTAGATGAAGATGGGTTGATCGCCt aataaCAGAGTTAGTGGTGAAGATATTATTATTGGTAAAACAACTCCTATACC TGAAGATGGGTTGATCGCCCCT |
| CaVip-13 | GCTACGACACCGTC ACCATT | CAATTAAACGAAGTGATATTAAAAATAGTGTCGCTACGACACCGTCACCAta ataaAAACACATCTAGTAATCGAATTAATAAACCCATGATAACATCAAACAA TAGCTACGACACCGTCACCATT |
| CaVip-14 | GTTGGATAATTCAG GTCAAC | AAGAACAAAAGAGACAACAAGAAGACGCGTCAAGGTTGGATAATTCAGGT taataaGCAGCACCAGCAACAAAATAACTCAAACGTAGTTTCAAACGGATCAA TTCGTTGGATAATTCAGGTCAAC |
| CaVip-15 | AACCTCAACCCAAC CAACAG | GACCACCACAACAACAACCTCAGCAACCTCAGCAACCTCAACCCAACCAAt aataaTCCCTTTCGACCAGACGAGTCACGAGCTCGTCGTTTGAATCAACCACT TGAACCTCAACCCAACCAACAG |

| | | |
|---|---|---|
| CaVip-16 | AAGGGAAGCCTCTG TTGGTT | CTCCGCAACCACGACCACCACAACAACAACCTCAGCAACCTCAGCAACCTt aataaCCAACAGAGGCTTCCCTTTCGACCAGACGAGTCACGAGCTCGTCGTTT GAAAGGGAAGCCTCTGTTGGTT |
| CaVip-17 | GATCCTAAGAAACC AATTGG | AAGTTGATGGTATGTCTGGTATGTTTAATCCTGATCCTAAGAAACCAATTtaa taaCATTATTGCCCATTCATCAACAACTAGATTATCGTTTAAAAAGGGAAGA GGATCCTAAGAAACCAATTGG |
| CaVip-18 | TGATCCTAAGAAAC CAATTG | CTCAAGTTGATGGTATGTCTGGTATGTTTAATCCTGATCCTAAGAAACCAtaa taaTAACATTATTGCCCATTCATCAACAACTAGATTATCGTTTAAAAAGGGA ATGATCCTAAGAAACCAATTG |
| CaVip-19 | GCAATAATGTTACC CCCAAT | TTGTTGCTCAAGTTGATGGTATGTCTGGTATGTTTAATCCTGATCCTAAGtaat aaTGGGGGTAACATTATTGCCCATTCATCAACAACTAGATTATCGTTTAAAA GCAATAATGTTACCCCCAAT |
| CaVip-20 | TCACCACCACCCAT ATCAAT | TTAGAACAGGTAAATCACAATTATGTCACACGTTAGCCGTTACGTGTCAGta ataaTGATATGGGTGGTGGTGAAGGAAAATGTCTTTACATTGATACTGAAGGT ATCACCACCACCCATATCAAT |
| CaVip-21 | GATGTATAGCTAAC TGCAGA | ATGACTTACAAAAGACATTACAATTTTCAACAAATCCCGATTTGAGACTAta ataaGCAGTTAGCTATACATCCTTGTTTCAAGTTTAAACCACAGTTGTCAGAA AGATGTATAGCTAACTGCAGA |
| CaVip-22 | TGTTACTGAGGCAT TCGTTC | ATAAAAATGGTGGGTCTTCGAGAATTCCACATTTTGTTACTGAGGCATTCtaa taaTTCAATCAAGATGAATTATATTCCCGATCCTGACGATTACAAGTTTAGGT TGTTACTGAGGCATTCGTTC |
| CaVip-23 | ATTGGGGTGATCTT AAACCT | AAAGTGAACTTGAAGACCTTCGTAATGGATTTGATTGGGGTGATCTTAAAta ataaCTACTTGTTCAAAGGTTTGTCATTTTACGTGTGTGGGAATAACTTGAGT GATTGGGGTGATCTTAAACCT |
| CaVip-24 | ATGGGTTCAGAAAC CCAGAT | TATTGAAGAATGTACAGCTGAAGTATGAAATTGATGGGTTCAGAAACCCAta ataaAAAAGTCAAACCAGAGTATTTAGAAAAATTTGGTGAAAATTTAGATCT TGATGGGTTCAGAAACCCAGAT |
| CaVip-25 | TCTGGCGATGAGTT CCAATC | ATTTCAACACCTTCTTACCTCCACCTACTCCGCCTAATTTGATTAATGGAtaat aaTTGGAACTCATCGCCAGAACCACATTCTCCTTCCCCAGGAAGATTGCAAC TCTGGCGATGAGTTCCAATC |

| | | |
|---|---|---|
| CaVip-26 | ACTCCATTCAATGG TGACAG | TGCATTTATCACCTTTGAAAAAACAATTACCAAACACTCCCACAAAGCAAta ataaCACCATTGAATGGAGTCCAGTTATATCACCAAACTCAAAGCAACCATTA CACTCCATTCAATGGTGACAG |
| CaVip-27 | GTATTAGAAGAAAC GACAGG | AATCTACTTCCACTGAAATTGAAATTGTAACAACCAGTTCTACTAAAGTTtaa taaTGTCGTTTCTTCTAATACTGATTTGACTAGTGAACCAACAAATACCAGAG GTATTAGAAGAAACGACAGG |
| CaVip-28 | GAAGCATACAAATA TCCACT | ATGGTGATAAAGATATCTCAATTGATGTTGAGTTTGAAAAGTCAACCGTTta ataaTGGATATTTGTATGCTTCCAGAGTTATGCCAAGTCTCAATAAGGTCACA AGAAGCATACAAATATCCACT |
| CaVip-29 | TGTTGAGAAGGATG TGGCAG | TTGGTGCAAGATCAAATCAACAGCTGACTCCACAACAAAATGCTGCACCAta ataaGCCACATCCTTCTCAACAAGGTCAAGCACAAGCTCAACATAATTTCCAG ATGTTGAGAAGGATGTGGCAG |
| CaVip-30 | TCTGTTGGTTGAGG CTGCTT | CATCTGGACAAACAACACCAAACATGTCACAACCTCCTAGTGCTGGCACGta ataaGCAGCCTCAACCAACAGAGCAAATGCGTCAATTACAAGACAAGCAACA GCTCTGTTGGTTGAGGCTGCTT |
| CaVip-Null-1 | ATGCCCCTCGGGAC CCCATG | NA |
| CaVip-Null-2 | TTGGGATAGGGCAC AGTACA | NA |
| CaVip-Null-3 | GGGGACCCCGGAGA GTGCAT | NA |
